# Novel role of Lin28 signaling in regulation of mammalian PNS and CNS axon regeneration

**DOI:** 10.1101/281725

**Authors:** Xue-Wei Wang, Chang-Mei Liu, Philip A. Hall, Jing-Jing Jiang, Christopher D. Katchis, Sehwa Kang, Bryan C. Dong, Shuxin Li, Feng-Quan Zhou

**Author notes:** Correspondence to Feng-Quan Zhou, Room 291, The John G. Rangos Sr. Bldg., 855 North Wolfe Street, Departments of Orthopaedic Surgery and Neuroscience, Johns Hopkins University School of Medicine, Baltimore, MD 21205.

## Abstract

Several signaling molecules involved in cellular reprogramming have been shown to regulate mammalian axon regeneration. We hypothesized that reprogramming factors are key regulators of axon regeneration. Here we investigated the role of Lin28, an important reprogramming factor, in the regulation of axon regeneration. We found that Lin28a and Lin28b and their regulatory partners, let-7 microRNAs (miRNAs), were both necessary and sufficient in regulating mature sensory axon regeneration in vivo. More importantly, overexpression of either Lin28a or Lin28b in mature retinal ganglion cells (RGCs) promoted robust and sustained optic nerve regeneration. Additionally, combined overexpression of Lin28a and downregulation of PTEN in RGCs acted additively to promote optic nerve regeneration by reducing the backward turning of regenerating RGC axons. Our findings not only identified a novel molecule promoting optic nerve regeneration but also suggested that reprogramming factors may play vital roles in regulating axon regeneration in mammals.

## Introduction

Axon regeneration in the mammalian central nervous system (CNS) always has been a struggling challenge in biomedical studies. Patients suffering from spinal cord injury, traumatic brain injury, glaucoma, and various neurodegenerative diseases, including Parkinson’s disease, Alzheimer’s disease, amyotrophic lateral sclerosis and multiple sclerosis, would greatly benefit from the discovery and development of approaches that can successfully regenerate CNS axons. CNS neurons are terminally differentiated cells that have lost a variety of intrinsic abilities supporting axon growth during the process of maturation. Recent studies have identified Klf4 and c-Myc, two reprogramming factors that can work together with other cohorts to reprogram differentiated cells back to induced pluripotent stem cells (iPSCs), as important regulators of axon regeneration (Belin et al., 2015; Moore et al., 2009). SOX11, a member of the SOX family to which another reprogramming factor SOX2 belongs, has also been reported to promote axon regeneration in non-α-RGCs (Norsworthy et al., 2017). Thus, it is likely that by manipulating reprogramming factors, mature CNS neurons can regain the intrinsic abilities to support axon regeneration after nerve injury.

Lin28 RNA-binding protein, originally discovered in *C*. *elegans* as a heterochronic regulator of larval development (Ambros and Horvitz, 1984; Moss et al., 1997), has also been demonstrated to be capable of enhancing the efficiency of the reprogramming from human somatic cells to iPSCs (Yu et al., 2007). A previous study in plant reported that *PpCSP1*, a *Lin28* homologue in the moss Physcomitrella patens, can reprogram differentiated leaf cells to stem cells (Li et al., 2017), indicating that the cellular reprogramming function of Lin28 is evolutionarily conserved from plants to mammals. Recent studies have shown that Lin28a and Lin28b, two paralogs in vertebrates, play vital roles in many growth-associated functions, such as body size and puberty control (Shinoda et al., 2013; Zhu et al., 2010), glucose metabolism (Shinoda et al., 2013; Zhu et al., 2011) and tissue regeneration (Shyh-Chang et al., 2013). In zebrafish, the reactivation of Lin28 was found essential for Müller glia dedifferentiation and retinal regeneration after retinal injury (Ramachandran et al., 2010). Abnormal expression of Lin28a and Lin28b has also been linked with different kinds of tumors (Nguyen et al., 2014; Tu et al., 2015; Urbach et al., 2014; Viswanathan et al., 2009; Wang et al., 2015; West et al., 2009). To our knowledge, the roles of Lin28 signaling in postmitotic neurons were only explored in two studies from the same lab (Amen et al., 2017; Huang et al., 2012). Lin28 proteins mainly function through let-7 miRNA family dependent manners by posttranscriptionally blocking the biogenesis of mature let-7 miRNAs (Heo et al., 2008; Nam et al., 2011; Newman et al., 2008; Rybak et al., 2008; Thornton et al., 2012; Viswanathan et al., 2008). Nonetheless, some let-7 independent mechanisms have also been discovered recently (Balzer et al., 2010; Shyh-Chang et al., 2013; Zeng et al., 2016). In *C*. *elegans*, let-7 has been reported to negatively regulate axon regeneration in anterior ventral microtubule neurons (Zou et al., 2013). In mammals, although glial let-7 miRNAs were shown to inhibit axon regeneration by targeting nerve growth factor in Schwann cells (Li et al., 2015), the function of neuronal let-7 miRNAs in axon regeneration remains unclear.

In our study, we found that Lin28a and Lin28b upregulation was rapidly induced in neurons of peripheral nervous system (PNS) following axon injury to support axon regeneration through subsequent downregulation of let-7 miRNAs. As a result, downregulation of Lin28a and Lin28b or upregulation of let-7a resulted in reduced sensory axon regeneration in vivo. Conversely, overexpression of Lin28a or Lin28b, or inhibition of let-7 miRNAs enhanced PNS sensory axon regeneration in vivo. Most importantly, we showed that overexpression of Lin28a or Lin28b produced robust and persistent CNS axon regeneration after optic nerve injury. In addition, we found that upregulation of Lin28a led to Akt activation and GSK3β inactivation in mature sensory neurons, as well as increased Ser235/236 phosphorylation of ribosomal protein S6 (pS6) in both sensory neurons and RGCs. Lastly, combined upregulation of Lin28a and downregulation of PTEN had an additive promoting effect on optic nerve regeneration, potentially through reducing the backward turning of regenerating axons. Our findings not only revealed Lin28 proteins as novel positive regulators of mammalian PNS/CNS axon regeneration, but also suggested that manipulation of reprogramming factors might be an effective way to promote axon regeneration in mature CNS neurons.

## Results

### Upregulation of Lin28a and Lin28b is necessary for sensory axon regeneration in vitro and in vivo

We first examined the expression of Lin28a and Lin28b in mouse DRGs during development. Lumbar 4 and 5 (L4/5) DRGs were dissected out from mice at ages of embryonic day 15 (E15), E18, postnatal day 0 (P0), P3, P7, P14, P21, P28 and P56, after which total RNA was extracted. Using qPCR, we found that the mRNA levels of Lin28a and Lin28b dropped sharply from E15 to birth, and remained relatively low through adulthood (Fig. S1A, B), indicating that Lin28a and Lin28b may be important in regulating axon growth of sensory neurons during early development. We next investigated if Lin28a and Lin28b are required in axon regeneration of adult sensory neurons. Sciatic nerve axotomy or sham surgery was performed on both sides of mice. Seven days after the surgery, L4/5 DRGs of each mouse were collected and total RNA was extracted. The levels of Lin28a and Lin28b mRNAs were significantly upregulated in L4/5 DRGs following sciatic nerve injury (SNI) (Fig. 1A). We also observed upregulated protein level of Lin28a 1 day or 3 days after SNI (Fig. 1B, C). The Lin28b antibody we used did not work well with mouse tissue extracts. These results suggested Lin28a and Lin28b might be positive regulators of sensory axon regeneration.

**Figure 1.**
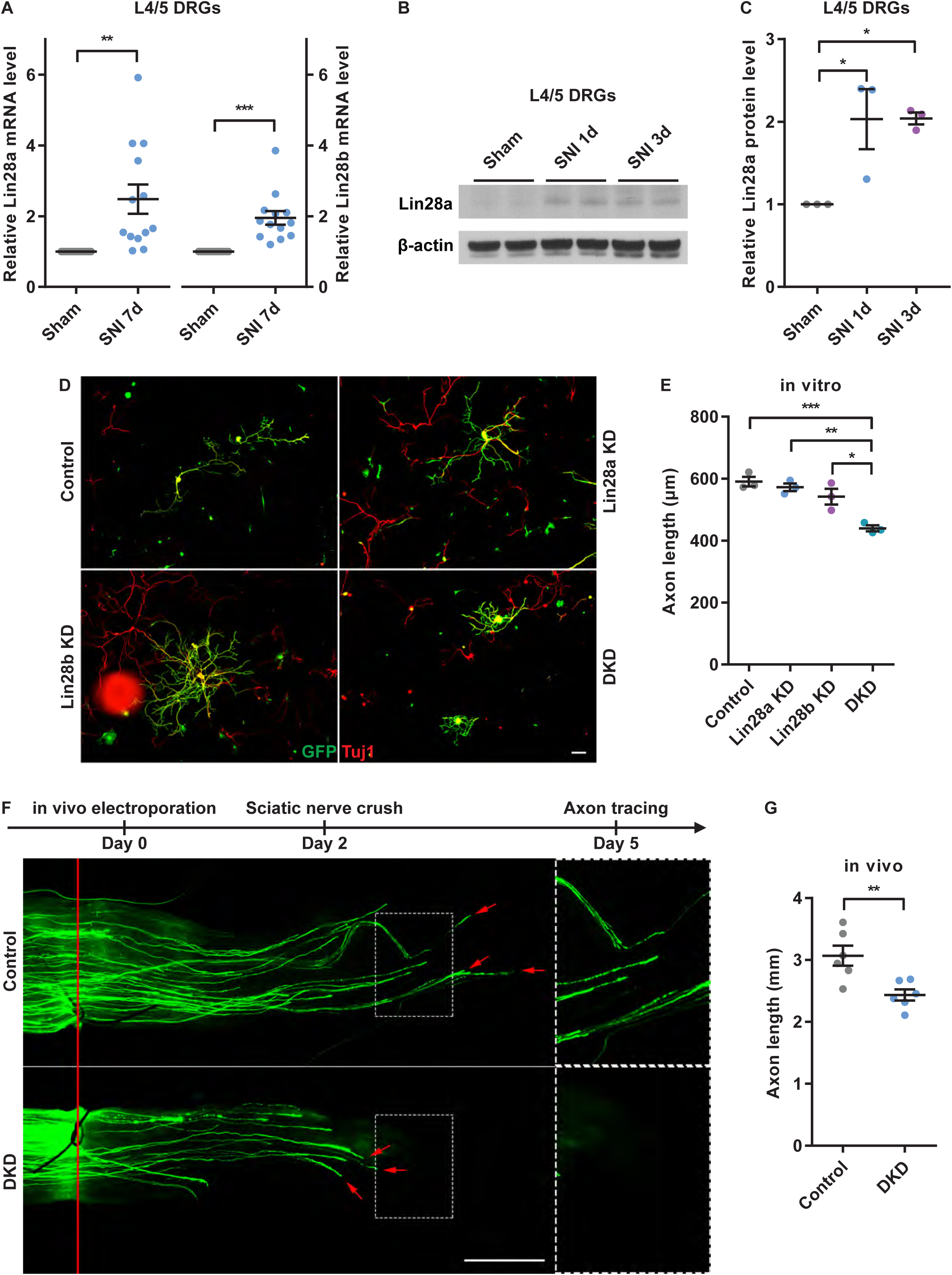
Upregulation of Lin28a and Lin28b was required for sensory axon regeneration in vitro and in vivo. (A) Lin28a (left) and Lin28b (right) mRNA levels were significantly increased in L4/5 DRGs 7 days after sciatic nerve injury (SNI) (one sample t test, *P* = 0.0039 for Lin28a, *P* = 0.0003 for Lin28b, n = 13 independent experiments). (B) Representative western blot results (2 out of 3 independent experiments) showing markedly increased Lin28a protein level in L4/5 DRGs 1 day or 3 days after SNI. (C) Quantification of relative Lin28a protein level in (B) (one-way ANOVA followed by Tukey’s multiple comparisons test, *P* = 0.0211, n = 3 independent experiments). (D) Representative images of cultured DRG neurons showing that simultaneously knocking down Lin28a and Lin28b impaired sensory axon regeneration in vitro. Cells were stained with anti-GFP (green) and anti-βIII tubulin (Tuj1, red) antibodies. Only axons of GFP^+^/Tuj1^+^ cells were measured. Scale bar, 100 µm. (E) Quantification of average axon length in (D) (one-way ANOVA followed by Tukey’s multiple comparisons test, *P* = 0.0009, n = 3 independent experiments). (F) Top: diagram showing the timeline of the experiment. Bottom: representative images of regenerating sciatic nerves showing that simultaneously knocking down Lin28a and Lin28b impaired sensory axon regeneration in vivo. The right column shows enlarged images of areas in the dashed white boxes in the left column. The red line indicates the crush sites. Red arrows indicate distal tips of regenerating axons. Scale bar, 1 mm for the left column, 0.5 mm for the right column. (G) Quantification of average axon length in (F) (unpaired Student’s t test, *P* = 0.0064, n = 6 mice in each group). \**P* < 0.05, \*\**P* < 0.01, \*\*\**P* < 0.001. Data are presented as mean ± SEM. KD, knockdown. DKD, double knockdown of Lin28a and Lin28b.

To further determine the functional roles of Lin28a and Lin28b in axon regeneration, we knocked down Lin28a and/or Lin28b by electroporating Lin28a siRNA (siLin28a) and/or Lin28b siRNA (siLin28b) together with GFP into dissociated adult DRG neurons and cultured them for 3 days. Electroporation of GFP only was used as control condition. The siLin28a or siLin28b was a group of four different siRNAs targeting different sequences of the same mRNA (ON-TARGETplus, Dharmacon). As a result, the concentration of each siRNA is relatively low, resulting in significantly reduced off-target effects. The knockdown efficacy of siLin28a or siLin28b was verified by detecting mRNA levels of Lin28a and/or Lin28b in cultured DRG neurons and proteins levels in a neuronal cell line, CAD cells, 3 days after the transfection of siRNAs. Results of both experiments demonstrated that the transfection of siLin28a and/or siLin28b successfully brought down the level of the corresponding mRNA (Fig. S2A) or protein (Fig. S2B). Interestingly, we found that the mRNA level of Lin28b significantly increased when Lin28a was knocked down by siLin28a in cultured DRG neurons (Fig. S2A), indicating that a compensatory upregulation of Lin28b mRNA level might occur when Lin28a is decreased. According to our previous study, the transfection efficiency of siRNAs using in vitro electroporation is over 95% in adult DRG neurons (Jiang et al., 2015), so nearly all GFP expressing neurons were also transfected with siRNAs. The results showed that compared to GFP transfection only, single knockdown of either Lin28a or Lin28b had no effect on sensory axon regeneration, whereas double knockdown of Lin28a and Lin28b significantly inhibited sensory axon regeneration in vitro (Fig. 1D, E), indicating that Lin28a and Lin28b play redundant roles in supporting sensory axon regeneration, resembling the discovery in the moss Physcomitrella patens that only quadruple deletion of all four *PpCSP* genes resulted in attenuated reprogramming (Li et al., 2017).

To extend the finding into in vivo model, we transfected GFP only or siLin28a and siLin28b along with GFP into left L4/5 DRGs of mice using in vivo electroporation, and the sciatic nerve crush was performed on the left side of each mouse 2 days later. Using the same in vivo electroporation approach, we have shown in a different study that sensory axon regeneration in vivo is slower during the first 2 days and reaches the fastest rate (∼1.5-2 mm/day) during the 3^rd^day after the sciatic nerve crush (see attached unpublished Table 1 in a different study). Therefore, we analyzed the inhibitory effect of Lin28a and Lin28b double knockdown 3 days after nerve injury when axons were in fast growth state. At the end of the third day, the left sciatic nerve of each mouse was collected and the lengths of all GFP labelled regenerating axons in the whole-mount nerve were measured. Over 96% of neurons in a DRG can be successfully transfected with siRNAs via in vivo electroporation (Fig. S2C, D), indicating that almost every GFP expressing sensory neuron was also transfected with siRNAs. The results showed that compared with the GFP only group, simultaneously knocking down Lin28a and Lin28b significantly impaired sensory axon regeneration in vivo (Fig. 1F, G), demonstrating that the axotomy-triggered upregulation of Lin28a and Lin28b is necessary for in vivo axon regeneration.

**Table 1.**
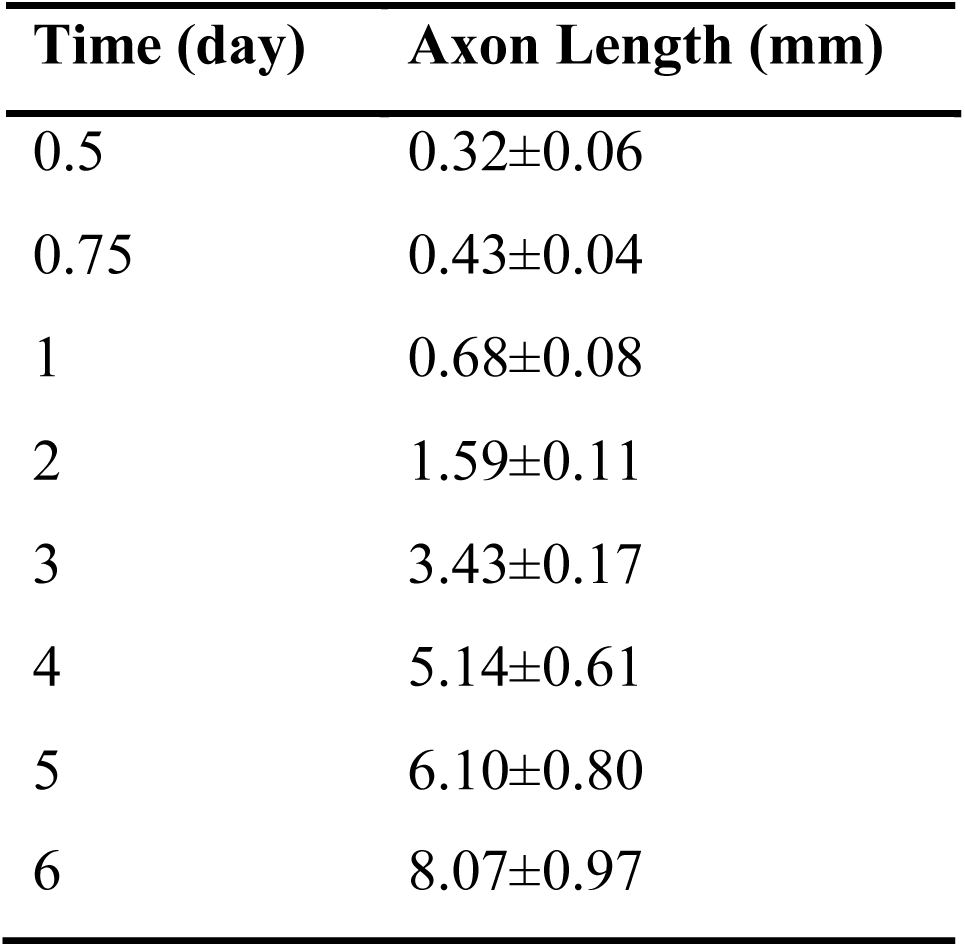
The length of regenerative axon in the sciatic nerve at the different time points. Axon lengths were presented as mean ± SD. This is a table from a different manuscript submitted elsewhere titled “Time course analysis of sensory axon regeneration in vivo by directly tracing regenerating axons in whole mount nerves”. This table is for reviewing purposes only. This is a partially brightened version of Figure 7A that is intended for better visualization for reviewing purposes only.

| Time (day) | Axon Length (mm) |
| --- | --- |
| 0.5 | 0.32±0.06 |
| 0.75 | 0.43±0.04 |
| 1 | 0.68±0.08 |
| 2 | 1.59±0.11 |
| 3 | 3.43±0.17 |
| 4 | 5.14±0.61 |
| 5 | 6.10±0.80 |
| 6 | 8.07±0.97 |

### Overexpression of Lin28a or Lin28b is sufficient to promote sensory axon regeneration in vivo

We then explored if overexpressing Lin28a or Lin28b sufficiently promotes sensory axon regeneration. Because single knockdown of either one of them had no effect on sensory axon regeneration, it indicated that either Lin28a or Lin28b alone is sufficient to support axon regeneration. We first used a mouse strain carrying a copy of human form *Lin28b* (hLin28b) that can be induced by a tetracycline transactivator *rtTA* located downstream of *Gt(Rosa26)Sor* promoter (hLin28b mice). Littermates with *rtRA* transgene only (M2rtTA mice) were used as control (for details, see Experimental Procedures). Following 2 days of tetracycline analog doxycycline (dox) treatment, high level of hLin28b was induced in L4/5 DRGs of hLin28b mice (Fig. 2A). On the third day of dox treatment, we electroporated GFP into L4/5 DRGs of M2rtTA mice and hLin28b mice to label the axons and after 2 days, the sciatic nerve crush was made. As mentioned above, within the first 2 days after nerve injury, sensory axons regenerate at slower rate. Thus, to evaluate the promoting effect of hLin28b upregulation, the nerves were harvested and the axons were measured 2 days after nerve injury. The results showed that the overexpression of hLin28b significantly enhanced sensory axon regeneration in vivo (Fig. 2B, C). To rule out the possibility that such promoting effect was caused by the overexpression of hLin28b in other cells in sciatic nerves, we constructed a CMV-Lin28a plasmid and specifically overexpressed Lin28a in L4/5 DRGs using in vivo electroporation (Fig. 2D). Control mice were electroporated with GFP. Two days after the sciatic nerve crush, the regenerating axons with overexpressed Lin28a were significantly longer than those in the GFP only group (Fig. 2E, F). These results demonstrated clearly that overexpressing either Lin28a or Lin28b sufficiently promotes sensory axon regeneration in vivo. Although the increase in axon length is only between 40% and 50%, it is considered to be very significant and robust because adult regenerating sensory neurons already have the fastest axon growth rate in any known model system.

**Figure 2.**
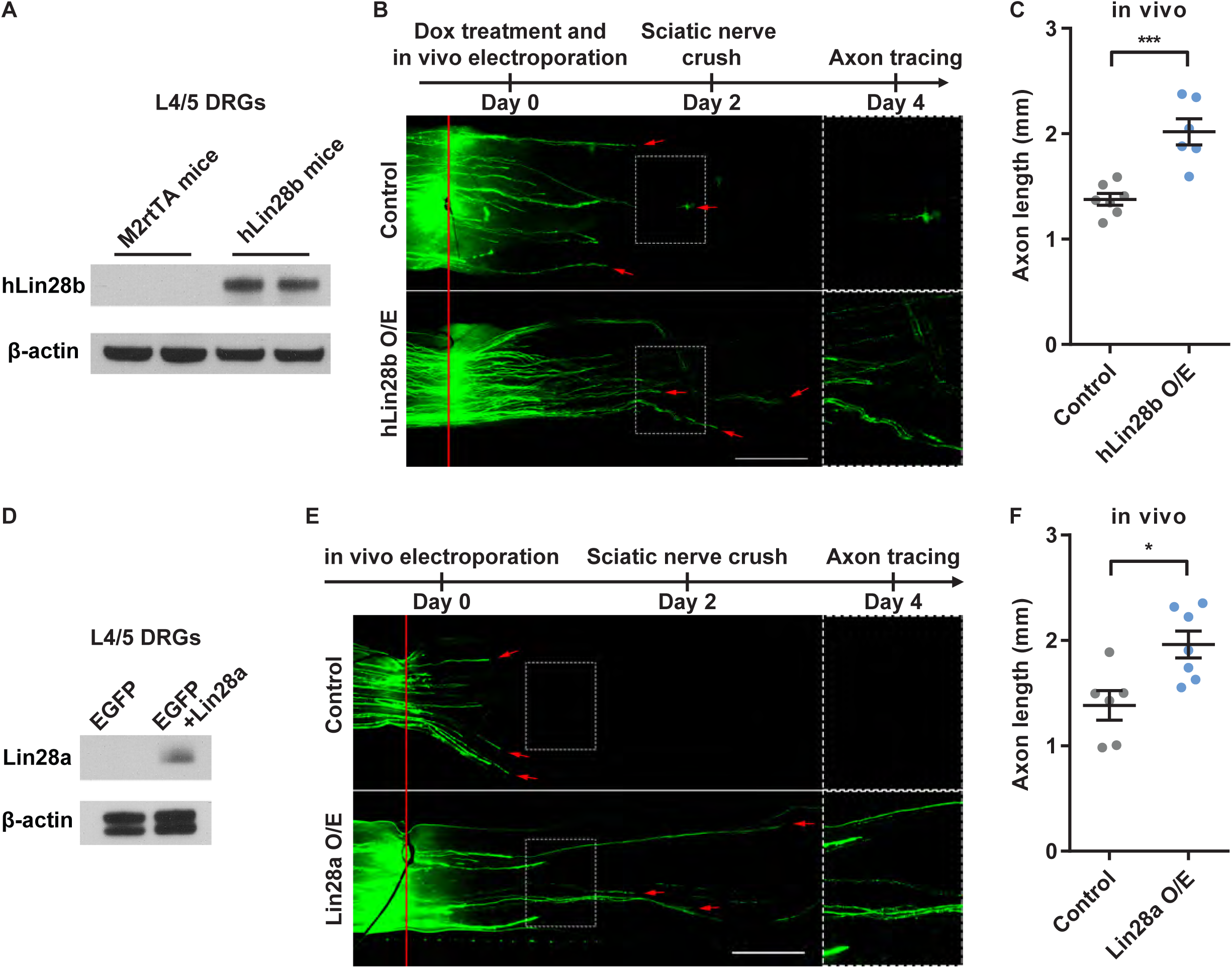
Overexpression of Lin28a or Lin28b sufficiently promoted sensory axon regeneration in vivo. (A) Representative western blot results (2 out of 3 independent experiments) showing markedly increased hLin28b protein level in L4/5 DRGs of hLin28b mice 2 days after induction of hLin28b expression by doxycycline. (B) Top: diagram showing the timeline of the experiment. Bottom: representative images of regenerating sciatic nerves showing that overexpression of hLin28b promoted sensory axon regeneration in vivo. The right column shows enlarged images of areas in the dashed white boxes in the left column. The red line indicates the crush sites. Red arrows indicate distal tips of regenerating axons. Scale bar, 1 mm for the left column, 0.5 mm for the right column. (C) Quantification of average axon length in (B) (unpaired Student’s t test, *P* = 0.0004, n = 7 and 6 mice in control group and hLin28b overexpression group, respectively). (D) Representative western blot result (1 out of 3 independent experiments) showing markedly increased Lin28a protein level in L4/5 DRGs 2 days after in vivo electroporation of CMV-Lin28a plasmid. (E) Top: diagram showing the timeline of the experiment. Bottom: representative images of regenerating sciatic nerves showing that overexpression of Lin28a promoted sensory axon regeneration in vivo. The right column shows enlarged images of areas in the dashed white boxes in the left column. The red line indicates the crush sites. Red arrows indicate distal tips of regenerating axons. Scale bar, 1 mm for the left column, 0.5 mm for the right column. (F) Quantification of average axon length in (E) (unpaired Student’s t test, *P* = 0.0109, n = 6 and 7 mice in control group and Lin28a overexpression group, respectively). \**P* < 0.05, \*\*\**P* < 0.001. Data are presented as mean ± SEM. O/E, overexpression.

### Downregulation of let-7 miRNAs is necessary for sensory axon regeneration in vitro and in vivo

As a renowned regulatory partner of Lin28 proteins, let-7 miRNA has been shown to be a negative regulator of axon regeneration in *C. elegans* (Zou et al., 2013). However, the role of mammalian let-7 miRNA family in axon regeneration has never been elucidated. We found that 7 days following SNI, the levels of let-7a and let-7b in L4/5 DRGs were significantly downregulated (Fig. 3A), indicating low levels of let-7 miRNAs in sensory neurons might be essential for sensory axon regeneration. To find out the functions of let-7 miRNAs in axon regeneration, we electroporated let-7a or let-7b mimic together with GFP into dissociated adult DRG neurons.Control neurons were only transfected with GFP. Data shown in Fig. 3B confirmed the overexpression of mature let-7a or let-7b in the cultured DRG neurons 3 days after electroporation. After culturing the DRG neurons for 3 days, we discovered that overexpression of either let-7a or let-7b drastically impaired sensory axon regeneration in vitro (Fig. 3C, E). Again, to confirm this finding in the in vivo model, we electroporated GFP only or let-7a mimic plus GFP into L4/5 DRGs 2 days before sciatic nerve crush and analyze the axon lengths 3 days after the crush. The result showed that compared with GFP only group, the overexpression of let-7a significantly reduced sensory axon regeneration in vivo (Fig. 3D, F). Mature let-7a level was confirmed to be dramatically elevated in L4/5 DRGs 2 days after in vivo electroporation of let-7a mimic (Fig. 3G). These results demonstrated that mammalian let-7 miRNAs are negative regulators of axon regeneration.

**Figure 3.**
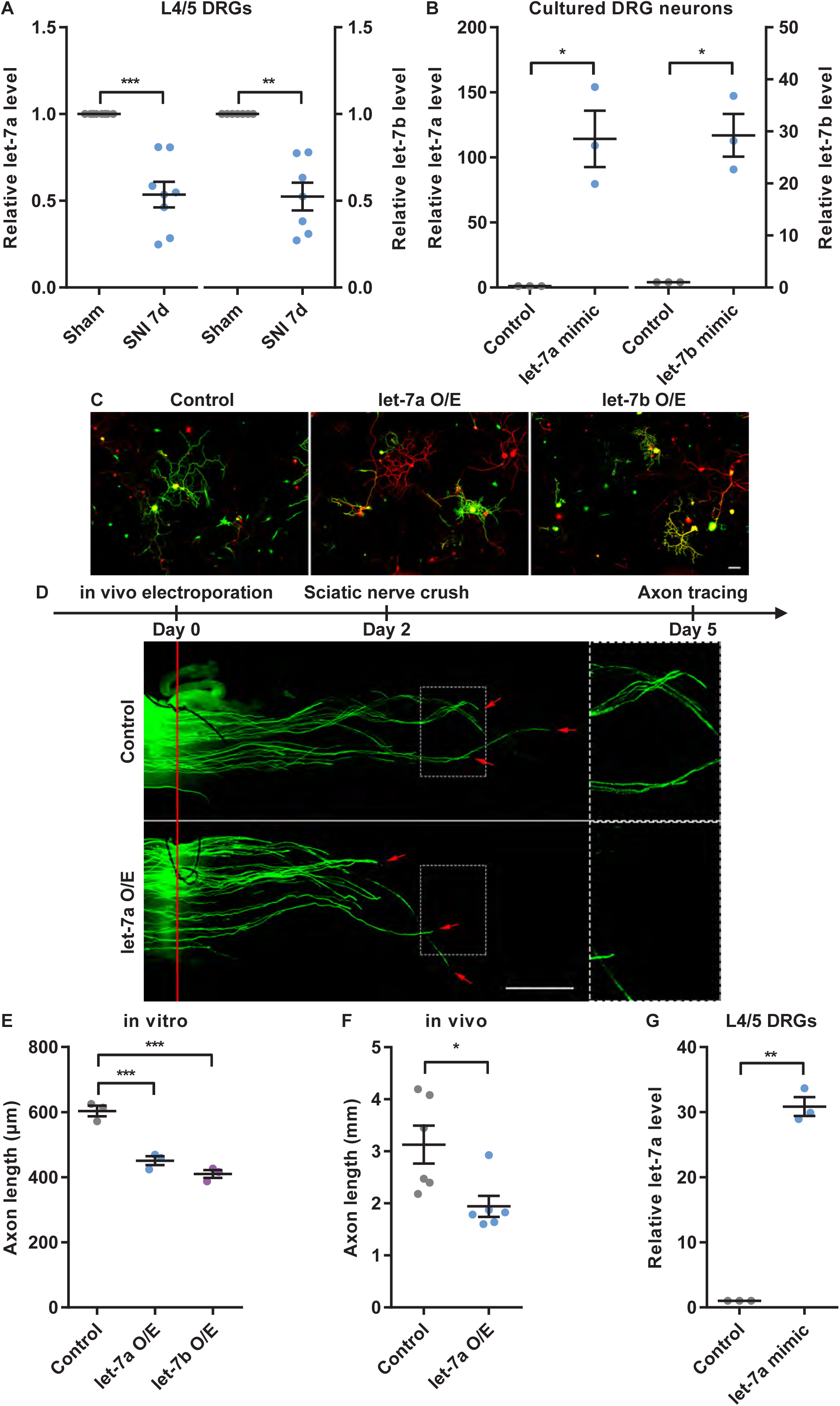
Downregulation of let-7 miRNAs was required for sensory axon regeneration in vitro and in vivo. (A) let-7a (left) and let-7b (right) levels were significantly decreased in L4/5 DRGs 7 days after sciatic nerve injury (SNI) (one sample t test, *P* = 0.0004 for let-7a, *P* = 0.0010 for let-7b, n = 8 independent experiments). (B) let-7a (left) or let-7b (right) levels in cultured DRG neurons were significantly increased 3 days after let-7a or let-7b mimic electroporation (one sample t test, *P* = 0.0346 for let-7a, *P* = 0.0205 for let-7b, n = 3 independent experiments). (C) Representative images of cultured DRG neurons showing that overexpression of let-7a or let-7b impaired sensory axon regeneration in vitro. Cells were stained with anti-GFP (green) and anti-βIII tubulin (Tuj1, red) antibodies. Only axons of GFP^+^/Tuj1^+^ cells were measured. Scale bar, 100µm. (D) Top: diagram showing the timeline of the experiment. Bottom: representative images of regenerating sciatic nerves showing that overexpression of let-7a impaired sensory axon regeneration in vivo. The right column shows enlarged images of areas in the dashed white boxes in the left column. The red line indicates the crush sites. Red arrows indicate distal tips of regenerating axons. Scale bar, 1 mm for the left column, 0.5 mm for the right column. (E) Quantification of average axon length in (C) (one-way ANOVA followed by Tukey’s multiple comparisons test, *P* = 0.0002, n = 3 independent experiments). Quantification of average axon length in (D) (unpaired Student’s t test, *P* = 0.0175, n = 6 mice in each group). (F) let-7a level in L4/5 DRGs was significantly increased 2 days after in vivo electroporation of let-7a mimic (one sample t test, *P* = 0.0023, n = 3 independent experiments). \**P* < 0.05, \*\**P* < 0.01, \*\*\**P* < 0.001. Data are presented as mean ± SEM. O/E, overexpression.

### let-7 miRNAs act downstream of Lin28a/b to regulate sensory axon regeneration in vivo

We next investigated the relationship between Lin28a/b and let-7 miRNA family in the regulation of axon regeneration. We harvested the L4/5 DRGs from hLin28b mice after 2 days of dox treatment and extracted the total RNA. Using qPCR, we found that the levels of almost all members of the let-7 miRNA family were significantly decreased compared with M2rtTA mice (Fig. 4A), consistent with the roles of Lin28a/b in suppressing mature let-7 biogenesis. Since Lin28 proteins and let-7 miRNAs usually respond to each other and form a double negative feedback loop (Viswanathan and Daley, 2010), we examined which one of them responds first to the axon injury signal and causes the subsequent change of the other one. We collected L4/5 DRGs at 0, 3, 6, 12, 24 hr after sciatic nerve axotomy and tested the levels of Lin28a and Lin28b mRNAs as well as the levels of let-7a and let-7b. The results showed that Lin28b started to escalate as early as 12.hr post-SNI, and both Lin28a and Lin28b mRNAs were significantly increased 24 hr post-SNI (Fig. 4B). In contrast, let-7a and let-7b remained at the baseline level even after 24 hr (Fig. 4B), indicating that SNI first induced the upregulation of Lin28a/b, which consequently led to the subsequent downregulation of let-7 miRNAs. Thus, we think that Lin28a/b upregulation acts upstream of let-7 downregulation in adult sensory neurons to regulate axon regeneration in response to nerve injury. To test this hypothesis, we overexpressed let-7a in L4/5 DRGs of dox-treated M2rtTA or hLin28b mice via in vivo electroporation and crushed the sciatic nerves 2 days later. Sensory axon regeneration in these mice was analyzed 2 days after nerve injury. The results showed that let-7a overexpression significantly blocked sensory axon regeneration promoted by hLin28b overexpression (Fig. 4C, D). Furthermore, we tested if direct inhibition of let-7 miRNAs sufficiently promotes sensory axon regeneration. Using in vivo electroporation of let-7 miRNA family inhibitor in L4/5 DRGs, levels of all let-7 miRNAs except miR-98 were significantly knocked down (Fig. 4E). The let-7 miRNA family inhibitor is a group of antisense oligonucleotides that have perfect sequence complementarity to all let-7 miRNAs, silencing all let-7 miRNAs with similar high efficiency. We found that the sensory axon regeneration in sciatic nerves was boosted to a degree comparable with hLin28b overexpression, indicating that knockdown of let-7 miRNAs in L4/5 DRGs can phenocopy hLin28b overexpression (Fig. 4C, D). These results provided strong evidence that Lin28a/b support sensory axon regeneration through their inhibition of let-7 miRNA family.

**Figure 4.**
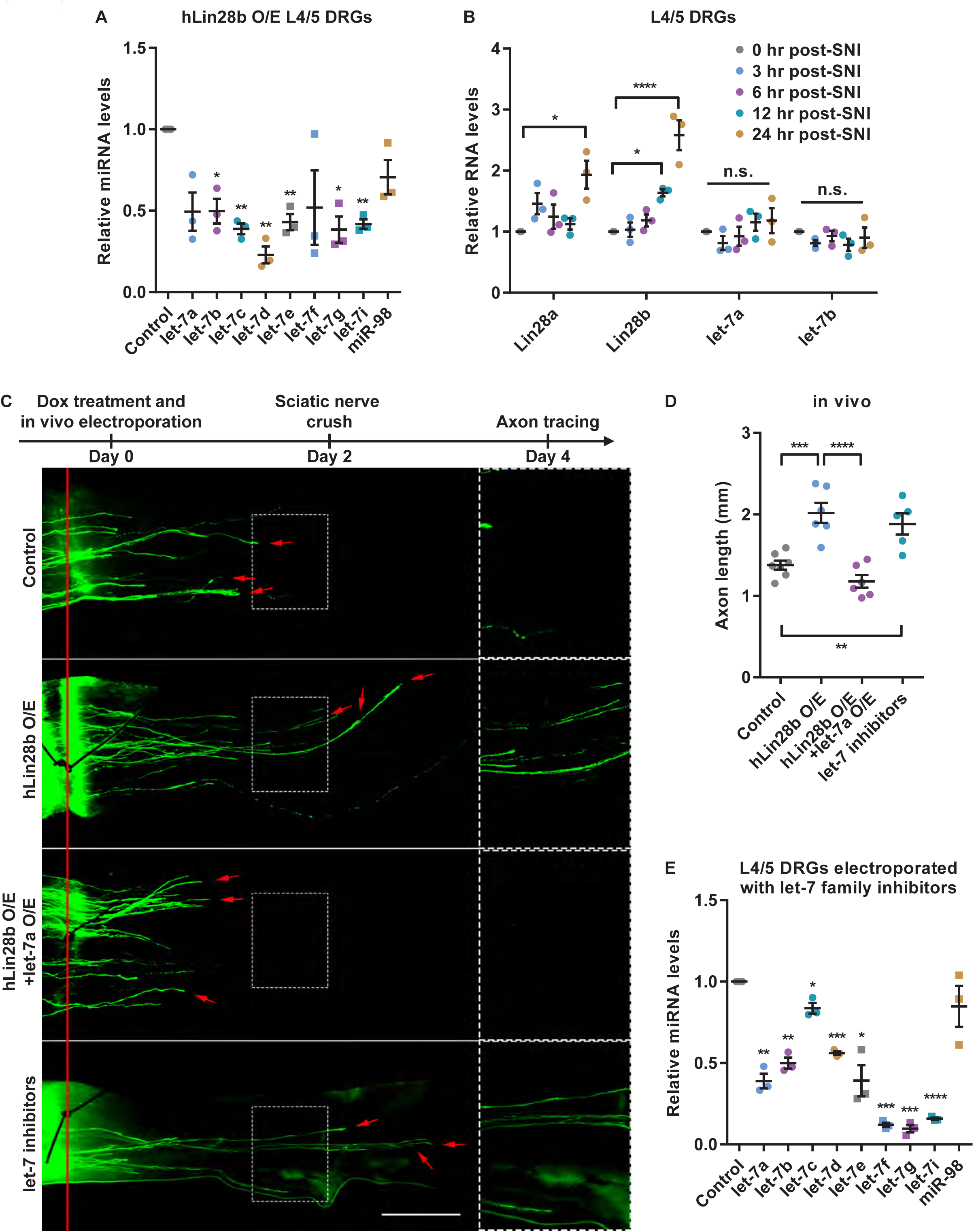
let-7 miRNAs acted downstream of Lin28 in regulating sensory axon regeneration in vivo. (A) Levels of let-7 miRNAs in L4/5 DRGs of hLin28b mice were significantly decreased 2 days after induction of hLin28b expression by doxycycline (one sample t test, *P* = 0.0501, 0.0221, 0.0029, 0.0045, 0.0074, 0.1702, 0.0167, 0.0124 and 0.1090 for let-7a, let-7b, let-7c, let-7d, let-7e, let-7f, let-7g, let-7i and miR-98, respectively, n = 3 independent experiments). (B) The increase of Lin28a and Lin28b mRNA levels occurred within 12-24 hr after sciatic nerve injury (SNI), while no significant change in let-7a or let-7b was found at 24 hr after SNI (one-way ANOVA followed by Tukey’s multiple comparisons test, *P* = 0.0164 for Lin28a, *P* < 0.0001 for Lin28b, *P* = 0.3634 for let-7a, *P* = 0.5388 for let-7b, n = 3 independent experiments). (C) Top: diagram showing the timeline of the experiment. Bottom: representative images of regenerating sciatic nerves showing that overexpressing let-7a abolished the enhanced sensory axon regeneration induced by hLin28b overexpression in vivo, and knocking down let-7 miRNAs was sufficient to promote sensory axon regeneration in vivo. The right column shows enlarged images of areas in the dashed white boxes in the left column. The red line indicates the crush sites. Red arrows indicate distal tips of regenerating axons. Scale bar, 1 mm for the left column, 0.5 mm for the right column. (D) Quantification of average axon length in (C) (one-way ANOVA followed by Tukey’s multiple comparisons test, *P* < 0.0001, n = 7 mice in control group, n = 5 mice in let-7 inhibitors group, n= 6 mice in other two groups). Note that the control group and the hLin28b overexpression group are identical to those shown in Fig. 2F. (E) Levels of let-7 miRNAs in L4/5 DRGs were significantly decreased 2 days after in vivo electroporation of let-7 miRNA family inhibitors (one sample t test, *P* = 0.0055, 0.0046, 0.0386, 0.0007, 0.0238, 0.0002, 0.0006, and 0.3509 for let-7a, let-7b, let-7c, let-7d, let-7e, let-7f, let-7g and miR-98, respectively, *P* < 0.0001 for let-7i, n = 3 independent experiments). n.s., not significant. \**P* < 0.05, \*\**P* < 0.01, \*\*\**P* < 0.001, \*\*\*\**P* < 0.0001, compared to control if not designated. Data are presented as mean ± SEM. O/E, overexpression.

### Upregulation of Lin28a in RGCs is sufficient to promote robust and sustained optic nerve regeneration

In light of the promising promoting effect of Lin28a/b on PNS sensory axon regeneration, we hypothesized that the overexpression of them might be able to enhance CNS axon regeneration as well. Therefore, we introduced the optic nerve regeneration model into our study. We started with the M2rtTA and hLin28b mice, in which optic nerve crushes were made on the right side on the same day when dox treatment started. Since over half of the hLin28b mice would die of hypoglycemia on the 10^th^ day after hLin28b induction, we assessed optic nerve regeneration 9 days after the injury. In fact, only one hLin28b mouse survived for 2 weeks in our experiment (Fig. S3A). The RGC axons were anterogradely labeled with Alexa 594-conjugated CTB, which was injected into the vitreous humor 2 days prior to tissue harvest. The fixed whole mount nerves underwent a tissue clearing procedure and were then imaged with confocal microscopy. Only very limited number of axons were able to cross the crush site in the control group 9 days post-injury (Fig. S3A). On the contrary, overexpression of hLin28b induced evident optic nerve regeneration (Fig. S3A, B). Induction of hLin28b in retinas of hLin28b mice was confirmed after 2 days of dox treatment (Fig. S3C). In order to exclude the likelihood that such an effect was the result of global overexpression of hLin28b and to investigate if stronger regeneration could be stimulated by extending the length of time and keeping the mice in a healthy state without metabolic disorder, we injected AAV-GFP or AAV2-Lin28a-Flag into the vitreous humors of mice and performed optic nerve crush 2 weeks after the viral injection. Two weeks after nerve crush, the nerves were collected and axon regeneration was evaluated. The virus transduction rate was about 88%, which was calculated by number of Flag^+^ RGCs/number of total RGCs in ganglion cell layer of whole-mount retinas (Fig. S4A, B), indicating that most RGCs were overexpressed with Lin28a. We found that compared with the GFP group, robust optic nerve regeneration was produced by the overexpression of Lin28a (Fig 5A, B). To further explore if overexpression of Lin28a can induce sustainable optic nerve regeneration in a longer timescale, we extended the regeneration time (the time between optic nerve crush and harvest) to 4 weeks. The result showed that overexpression of Lin28a significantly increased not only the length but also the number of regenerating axons compared to the 2-week group (Fig. 5A, B). Assessment of RGC survival showed no difference between the GFP group and the Lin28a overexpression group either 2 weeks or 4 weeks after optic nerve crush, but instead revealed a significantly lower RGC survival rate in the retinas 4 weeks after nerve crush compared to 2 weeks after nerve crush (Fig. S5A, B), indicating that the death of RGCs continued throughout the 4 weeks after optic nerve crush, and that Lin28a promoted optic nerve regeneration via boosting regeneration potential in neurons that survived the optic nerve crush rather than by protecting neurons from dying of the injury. We also estimated the shrinkage rate of the optic nerves caused by tissue clearing. We found that the nerves shrank to 82.0 ± 2.81% in length (Fig. S6A, B) and 56.3 ± 6.58% in diameter (Fig. S6A, C), resulting in a total shrinkage of 73.7 ± 5.72% in volume (estimated using the formula for the volume of a cylinder) (Fig. S6A, D). Thus, the axon length observed in our experiments was actually underestimated by about 18%. Overall, these results demonstrated that overexpression of hLin28b or Lin28a can induce robust and sustainable optic nerve regeneration without affecting the survival rate of RGCs.

**Figure 5.**
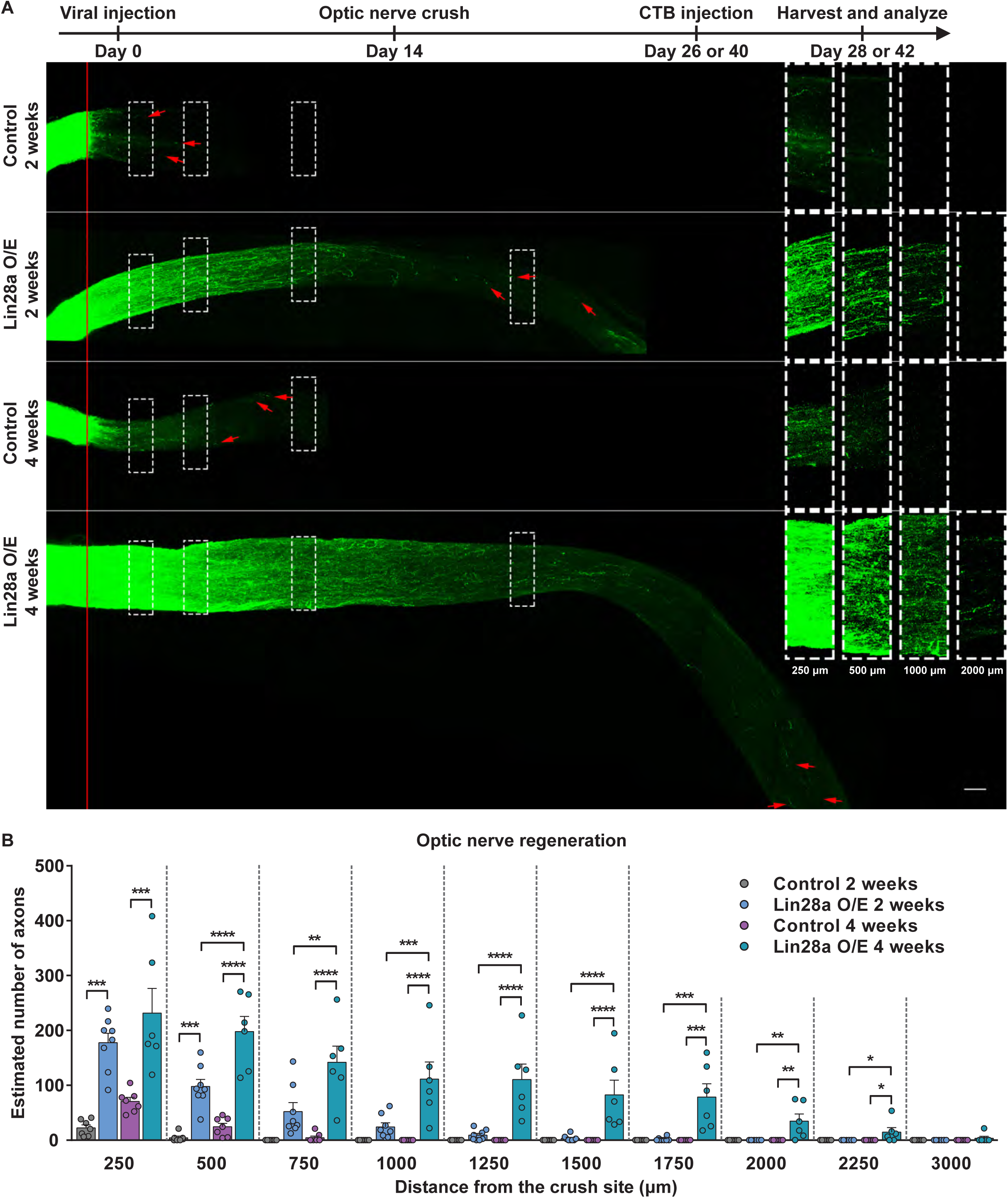
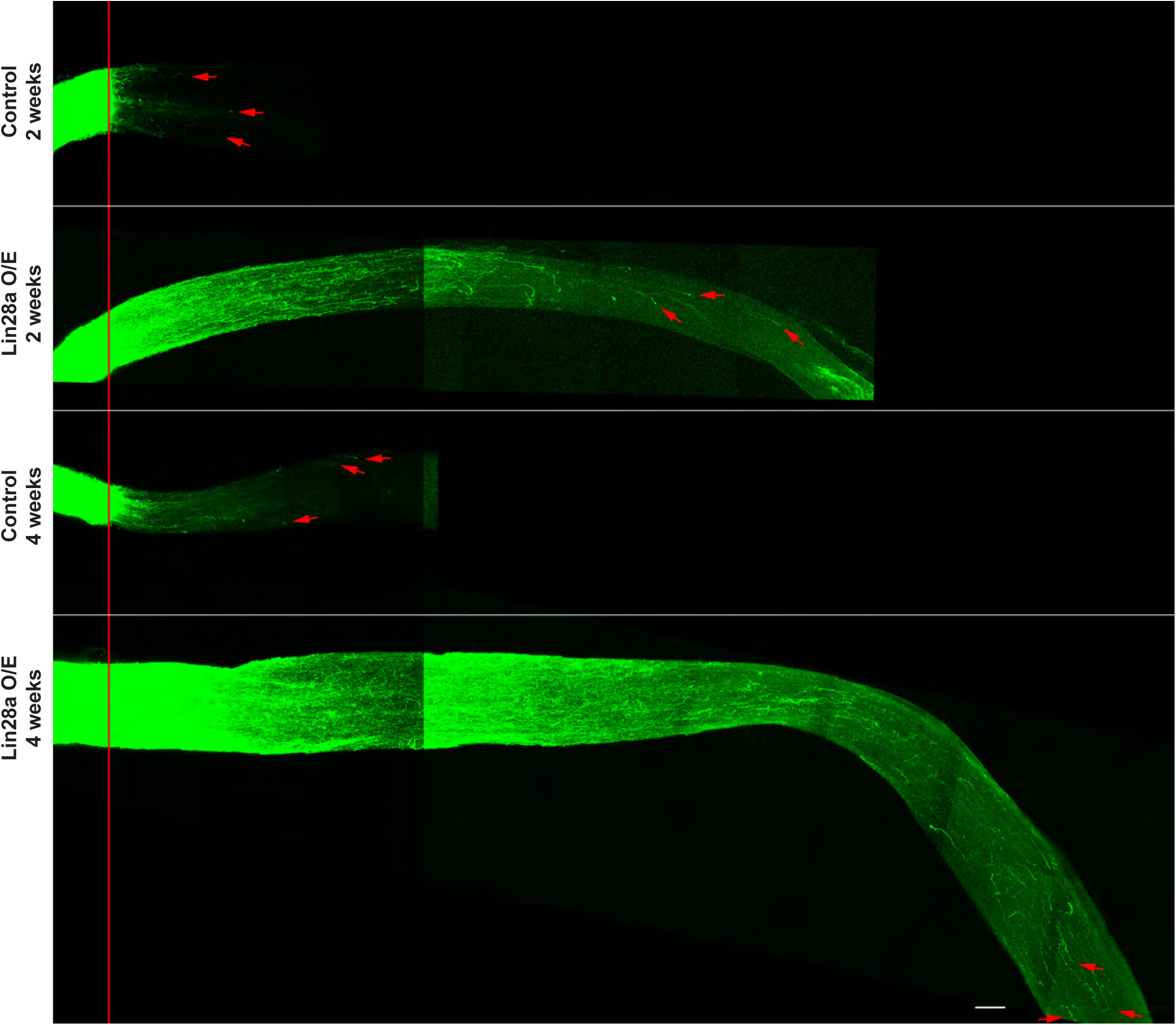
Overexpression of Lin28a in RGCs induced robust and sustained optic nerve regeneration. (A) Top: diagram showing the timeline of the experiment. Bottom: representative images of cleared whole-mount optic nerves showing that overexpression of Lin28a in RGCs induced drastic and persistent axon regeneration in optic nerves 2 and 4 weeks after nerve crush. The right column shows enlarged images of nerves at 250, 500, 1000 and 2000 µm distal to the crush sites, which are marked by dashed white boxes in the left column. The red line indicates the crush sites. Red arrow shows the longest axon of each nerve. Scale bar, 100 µm for the left column, 50 µm for the right column. O/E, overexpression. (B) Quantification of estimated number of axons at certain distances from the crush sites of optic nerves in (A) (one-way ANOVA followed by Tukey’s multiple comparisons test, *P* < 0.0001 at 250, 500, 750, 1000, 1250, 1500 and 1750 µm, *P* = 0.0005, 0.0193 and 0.3113 at 2000, 2250 and 3000 µm, respectively, n = 7 mice in 2-week and 4-week control groups, n = 8 mice in 2-week Lin28a overexpression group, n = 6 mice in 4-week Lin28a overexpression group). \**P* < 0.05, \*\**P* < 0.01, \*\*\**P* < 0.001, \*\*\*\**P* < 0.0001. Data are presented as mean ± SEM. O/E, overexpression.

### Lin28a/b regulate Akt, GSK3β, and mTOR signaling in sensory neurons and RGCs

Lin28/let-7 axis has previously been proven to regulate glucose metabolism through the insulin-PI3K-Akt pathway (Zhu et al., 2011). Additionally, increased level of pS6, downstream of PI3K-Akt-mTOR pathway, was linked to PTEN knockout-induced optic nerve regeneration (Park et al., 2008). GSK3β, the phosphorylation at Ser9 of which by Akt inhibits its activity, has been shown to negatively control axon regeneration (Guo et al., 2016; Leibinger et al., 2017). Thus, it is reasonable to speculate that the Lin28-induced PNS and CNS axon regeneration is associated with the regulation of Akt, GSK3β and/or mTOR signaling. We thus collected L4/5 DRGs from hLin28b mice and M2rtTA control mice after 2 days of dox treatment and examined the level of Ser473 phosphorylation of Akt (pAkt), Ser9 phosphorylation of GSK3β (pGSK3β), and pS6. We found that overexpression of hLin28b significantly increased the levels of pAkt, pGSK3β, and pS6 (Fig. 6A, B). Importantly, we also observed that 2 weeks after optic nerve crush, the percentage of pS6^+^ RGCs increased about three-fold in the Lin28a overexpression group compared to the GFP control group (Fig. 6C, D). These results suggested that Lin28/let-7 axis might mediate axon regeneration, at least partially, through Akt activation, GSK3β inactivation, and/or activation of the mTOR pathway.

**Figure 6.**
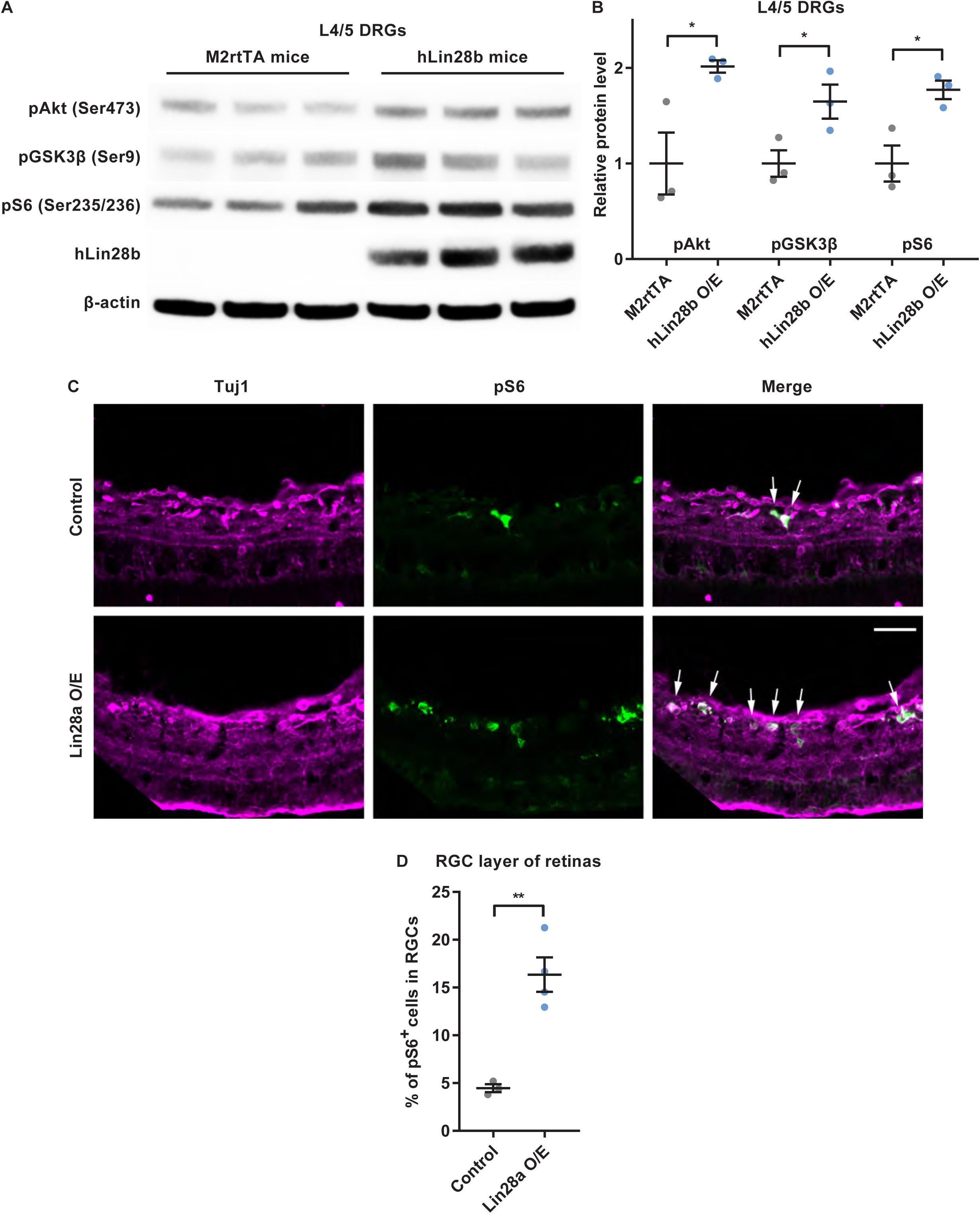
Lin28 overexpression in sensory neurons or RGCs was associated with Akt, GSK3β, and mTOR signaling. (A) Western blot results of 3 independent experiments showing overexpression of hLin28b elevated levels of pAkt, pGSK3β and pS6 in L4/5 DRGs. (B) Quantification of relative pAkt, pGSK3β and pS6 levels in (A) (unpaired Student’s t test, *P* = 0.0372, 0.0217 and 0.0453 for pAkt, pGSK3β and pS6, respectively, n = 3 independent experiments). (C) Representative images of retinal cyrosections showing that overexpression of Lin28a in retinas markedly increased the percentage of pS6^+^ RGCs in RGC layer. Cryosections were stained with anti-pS6 (green) and anti-βIII tubulin (Tuj1, far-red) antibodies. White arrows indicate pS6^+^ RGCs. Scale bar, 50 µm. (D) Quantification of percentage of pS6^+^ RGCs in RGC layer of control retinas and Lin28a overexpression retinas (unpaired Student’s t test, *P* = 0.0027, n = 3 mice in control group, n = 4 mice in Lin28a overexpression group). \**P* < 0.05, \*\**P* < 0.01. Data are presented as mean ± SEM. O/E, overexpression.

### Lin28a overexpression and PTEN knockdown have additive effect on optic nerve regeneration

We next tested if Lin28a overexpression and PTEN knockdown have additive effect on optic nerve regeneration. Viruses expressing shRNA against PTEN (AAV2-shPTEN) and AAV2-Lin28a-Flag were injected into the vitreous humors of the mice on day 0 and day 2 of the experiment, respectively, and the optic nerves were crushed 2 weeks after the injections. Control mice were injected with AAV2-GFP. Two weeks after the crush, we collected the nerves and found that combining Lin28a overexpression and PTEN knockdown induced faster optic nerve regeneration (Fig. 7A, B). Specifically, compared to Lin28a overexpression only and PTEN knockdown only, the combination of Lin28a overexpression and PTEN knockdown induced significantly more axons to regenerate to 750, 1000 and 1250 µm from the crush site. Additionally, at 1500, 1750 and 2000 µm from the crush site, where almost no axons could be observed in single treatment groups, combined Lin28a overexpression and PTEN knockdown caused some axons to reach these spots. (Fig. 7B). These results indicated that Lin28a overexpression and PTEN knockdown act additively in promoting optic nerve regeneration.

**Figure 7.**
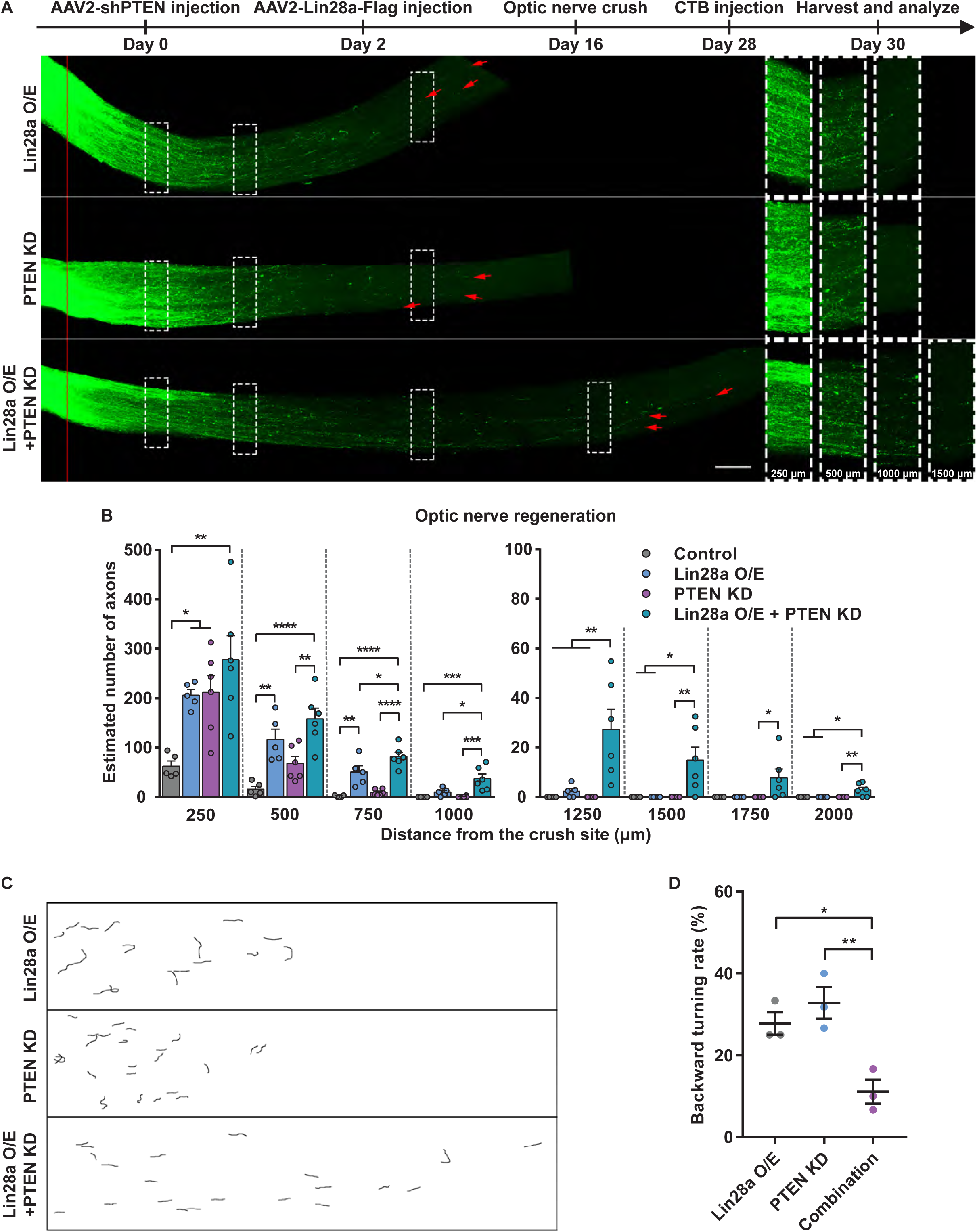
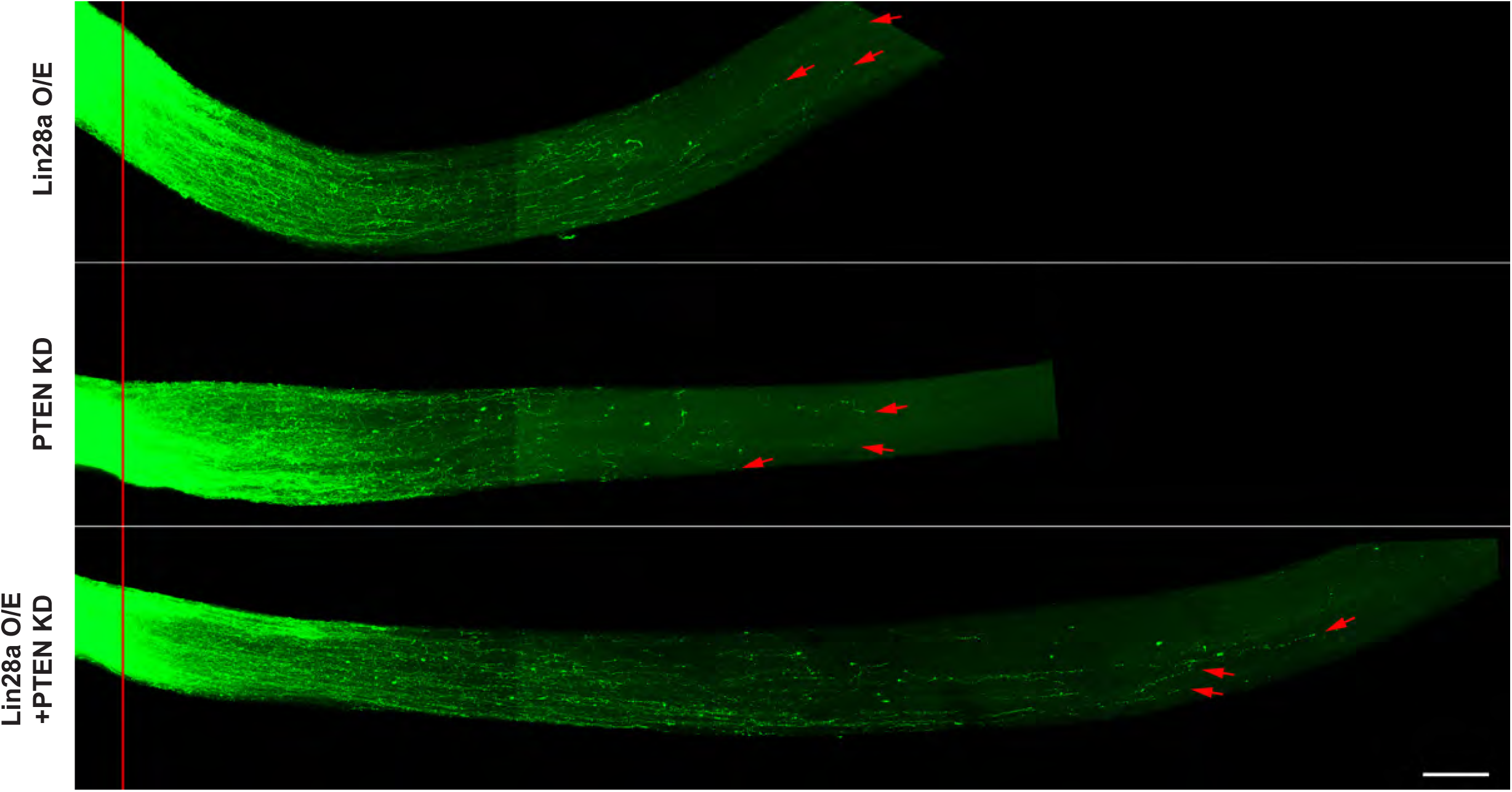
Combination of Lin28a overexpression and PTEN knockdown had additive effect on optic nerve regeneration. (A) Top: diagram showing the timeline of the experiment. Bottom: representative images of cleared whole-mount optic nerves showing that combined Lin28a overexpression and PTEN knockdown in RGCs induced faster axon regeneration in optic nerves 2 weeks after nerve crush. The right column shows enlarged images of nerves at 250, 500, 1000 and 1500 µm distal to the crush sites, which are marked by dashed white boxes in the left column. The red line indicates the crush sites. Red arrow shows the longest axon of each nerve. Scale bar, 100 µm for the left column, 50 µm for the right column. (B) Quantification of estimated number of axons at certain distances from the crush sites of optic nerves in (A) (one-way ANOVA followed by Tukey’s multiple comparisons test, *P* = 0.0025, 0.0001, 0.0003, 0.0006, 0.0028, 0.0242 and 0.0039 at 250, 500, 1000, 1250, 1500, 1750 and 2000µm, respectively, *P* < 0.0001 at 750 µm, n = 5 mice in control group and Lin28a overexpression group, n = 6 mice in PTEN knockdown group and combination group). (C) Representative reconstructed graphs of axon trajectories showing that the combination of Lin28a overexpression and PTEN knockdown significantly reduced the backward turning rate of axons. (D) Quantification of backward turning rate of axons in (C) (one-way ANOVA followed by Tukey’s multiple comparisons test, *P* = 0.0075, n = 3 mice in each group). \**P* < 0.05, \*\**P* < 0.01, \*\*\**P* < 0.001, \*\*\*\**P* < 0.0001. Data are presented as mean ± SEM. O/E, overexpression. KD, knockdown.

Previous studies have shown that although robust optic nerve regeneration can be achieved by stimulating intrinsic growth ability, long-distance regeneration and precise functional targeting are still largely hindered by axonal misguidance (Pernet et al., 2013; Pernet and Schwab, 2014; Yungher et al., 2015). Thus, we quantified the backward turning rate of regenerating optic nerve axons under each condition. The results showed about 30% of regenerating axons induced by either Lin28a overexpression or PTEN knockdown alone made backward turning (Fig. 7C, D). Combination of Lin28a overexpression and PTEN knockdown significantly reduced the percentage of axons that turned backwards (Fig. 7C, D). These results suggested that reduced backward turning rate of regenerating axons at least partially explains the additive effect of Lin28a overexpression and PTEN knockdown.

## Discussion

Throughout development, stem cells undergo many steps to turn into differentiated cells. During such processes, the whole gene expression profile changes drastically with stem cell related genes shut down and only genes relevant for the differentiated cell type expressed. The emerging idea is that the regulation of gene expression during cell differentiation is largely achieved through epigenetic modifications, including DNA methylation, histone modifications, and non-coding RNAs. Importantly, differentiated cells (e.g. fibroblast) can be reprogrammed back to iPSCs by overexpressing several reprogramming factors, such as Sox2, c-Myc, Klf4, Oct4, Nanog, and Lin28 (Takahashi and Yamanaka, 2006; Yu et al., 2007), which leads to global epigenetic remodeling. Do mammalian CNS neurons undergo similar epigenetic changes during maturation to lose their ability to support axon growth? If so, can we manipulate reprogramming factors to help CNS neurons reclaim such ability?

Indeed, Klf4 (Moore et al., 2009), c-Myc (Belin et al., 2015), and Sox11 (Norsworthy et al., 2017) are all important regulators of axon regeneration. A previous study reported that the reactivation of Lin28a, another reprogramming factor used in inducing iPSCs from human somatic cells, can promote mammalian tissue regeneration in postnatal tissues by reprogramming cellular metabolism (Shyh-Chang et al., 2013). These evidences led us to hypothesize that CNS neurons can also regain the potential to support axon regeneration by reintroduction of Lin28 proteins. Our study showed clearly that Lin28a and Lin28b not only are required, at least partially, for intrinsic axon regeneration ability in PNS sensory neurons, but also can promote both PNS and more importantly CNS axon regeneration in vivo when overexpressed. Our findings support the notion that mature CNS neurons can restore axon regeneration ability by modulating reprogramming factors. However, further studies are required to reveal if and how Lin28 signaling reprograms gene expression or metabolic state in mature mammalian neurons (Fig. S6E). For instance, a recent study showed that during iPSC reprogramming Lin28 acted to regulate mitochondrial function, nucleotide metabolism, and histone methylation (Zhang et al., 2016). Another study showed that Lin28a functioned to recruit 5-methylcytosine-diocxygenase Tet1 to regulate DNA methylation and gene expression (Zeng et al., 2016). Moreover, since cell reprogramming can now be achieved by using small-molecule compounds (Hou et al., 2013; Zhao et al., 2015), an easy-to-manage method without genomic integration, new translational opportunities of enhancing CNS axon regeneration may be opened.

The major downstream signaling mediator of Lin28 is let-7 miRNA family, which has been broadly studied due to its phylogenetically conserved expression patterns and functions. It has been reported previously in *C. elegans* that let-7 inhibits axon regeneration in older AVM neurons, whereas its loss of function reactivates AVM axon regeneration (Zou et al., 2013). In our study, we found that let-7 gain of function inhibited axon regeneration and abrogated the hLin28b overexpression-induced acceleration of axon regeneration in injured mouse sciatic nerves, while let-7 loss of function significantly improved axon regeneration, indicating that the role of let-7 as a negative regulator in axon regeneration is also evolutionarily conserved between nematodes and mammals. However, due to gene duplication, various members exist in the let-7 miRNA family in mammals, making the situation a lot more complicated than it is in worms. Distinct roles played by different let-7 miRNAs in the regulation of mammalian axon regeneration are yet to be elucidated in future studies.

Here we found that Lin28a upregulation led to Akt activation, GSK3β inactivation, and mTOR activation in mature sensory neurons and RGCs. These results suggest that the Lin28-promoted optic nerve regeneration might be partially mediated by the Akt-mTOR and/or GSK3β.pathways. Future studies using rapamycin or active mutant of GSK3β are required to determine if these pathways functionally act downstream of Lin28 to promote axon regeneration. We showed that Lin28a overexpression and PTEN knockdown acted additively to promote optic nerve regeneration, indicating that these two approaches also have some different underlying mechanisms although they both regulate the Akt-mTOR and GSK3β pathways.

Tissue clearing techniques have been widely adopted in recent studies due to their irreplaceable advantages in imaging experiments. Different clearing procedures, however, may yield different tissue shrinkage rate, resulting in inconsistencies in studies involving measurement of tissue size in various dimensions. Our observation suggested that the tissue shrinkage rate must not be overlooked and should be taken into consideration when comparing results from different tissue clearing techniques.

## Acknowledgements

We thank Dr. George Daley and Dr. Angelika Doetzlhofer for sharing mouse strains. The study was supported by grants (to F.-Q.Z. and S.L.) from NIH (R01NS064288, R01NS085176, R01GM111514, R01EY027347, 1R01NS079432, 1R01EY024575), the Craig H. Neilson Foundation, the BrightFocus Foundation and Shriners Research Foundation (SHC-86300-PHI, SHC-86200-PHI-16 and SHC-85100).

## Author Contributions

F.-Q.Z., X.-W.W., C.-M.L., and J.-J.J. initiated the projects and designed the experiments. F.-Q.Z., X.-W.W., and S.L. discussed about the experimental design. X.-W.W. performed experiments and analyzed data. P.A.H., C.D.K., S.K., and B.C.D. helped analyze imaging data and genotype mice. X.-W.W. and F.-Q.Z. wrote the paper. B.C.D. and S.L. helped proofread the manuscript.

## Declaration of Interests

The authors declare no competing interests.

## Experimental Procedures

### Animals

All animal experiments were completed according to the animal protocol approved by the Institutional Animal Care and Use Committee of the Johns Hopkins University. Female CF-1 mice of 6-8 weeks old were purchased from Charles River Laboratories. The *hLin28b* mutant mouse strain (JAX stock #023911) (Zhu et al., 2011) was a kind gift from Dr. George Daley’s laboratory at Harvard Medical School. The *M2rtTA* transgenic line (JAX stock #006965) (Hochedlinger et al., 2005) was obtained from Dr. Angelika Doetzlhofer’s laboratory at Johns Hopkins University School of Medicine. Male *M2rtTA*^tg/tg^ mice were crossed with female *hLin28b*^tg/+^ mice to generate *M2rtTA*^tg/+^; *hLin28b*^tg/+^ mice and *M2rtTA*^tg/+^ mice, which were used as hLin28b mice and M2rtTA control mice in experiments, respectively. Mice of both sexes were used. Genotypes of the mice were determined by PCR and sequences of genotyping primers were from The Jackson Laboratory. Doxycycline-containing food (Bio-Serv F3893) was used to induce the expression of *hLin28b*. All animal surgeries were performed under anesthesia induced by intraperitoneal injection of ketamine (100 mg/kg) and xylazine (10 mg/kg) diluted in sterile saline.

### Constructs

pCMV-GFP was a gift from Dr. Connie Cepko (Addgene plasmid # 11153). pMSCV-mLin28A was a gift from Dr. George Daley (Addgene plasmid # 26357). The Lin28a open reading frame was PCR-amplified from pMSCV-mLin28A using a forward primer that incorporated a 5’ EcoRI restriction site (5’-CGGAATTCATGGGCTCGGTGTCCAACC-3’) and a reverse primer that incorporated a 3’ NotI restriction site (5’-ATAAGAATGCGGCCGCTCAATTCTGGGCTTCTGGG-3’). The amplified sequence was then used to replace the EGFP open reading frame in pCMV-GFP with standard digestion and ligation. The Lin28a-Flag open reading frame with a 5’ BamHI restriction site and a 3’ EcoRV restriction site was synthesized using gBlocks (codon optimized) and was used to replace EYFP open reading frame in pAAV-EF1a-EYFP. The shPTEN plasmid was a gift from Drs. David Turner and Kevin Park. All enzymes were purchased from New England Biolabs.

### Western blot analysis

Proteins were extracted from DRG or retinal tissues, or CAD cells using the RIPA buffer containing protease inhibitor cocktail and phosphatase inhibitor cocktail. The extracted proteins were then separated by 4-12% gradient SDS-PAGE gel electrophoresis and transferred onto polyvinylidene fluoride membranes. After being blocked with 5% non-fat milk, blots were incubated overnight with primary antibodies against target proteins at 4°C, followed by corresponding HRP-conjugated secondary antibodies for 1 hr at room temperature. Blots were washed with 1x TBS containing 0.1% tween-20 for 15 min for 4 times after each antibody incubation. Bands from 3 independent experiments were analyzed with the ImageJ software (NIH). Band density was first normalized to the loading control, β-actin, and then normalized to the control group. Antibodies against Lin28a (1:1000, 3978S), Lin28b (1:1000, 5422S), hLin28b (1:1000, 4196S), pS6 Ser235/236 (1:2000, 4858S), pAkt Ser473 (1:2000, 4060S), pGSK3β Ser9.(1:1000, 5558S), c-Myc (1:1000, 5605S) were purchased from Cell Signaling Technology. Anti-.β-actin antibody (1:10000, A1978) was from Sigma-Aldrich.

### Quantitative real-time polymerase chain reaction (qPCR)

Total RNA from cells or tissues was first isolated with the miRNeasy mini kit (Qiagen), and then reversely transcribed to first strand cDNA using Transcriptor First Strand cDNA Synthesis Kit (Roche). For mRNAs, anchored oligo(dT)_18_ primer was used, while for miRNAs, gene-specific stem-loop primers were used following procedures described in a previous study (Chen et al., 2005). To detect the level of mRNAs or miRNAs, 10 ng first strand cDNA was amplified with gene-specific primers and LightCycler 480 SYBR Green I Master (Roche) using the LightCycler 480 II (Roche). All experiments were done in triplicate. Relative levels of mRNAs or miRNAs were determined using the ddCt method and normalized to the control group. Gapdh and Rnu6b were used as the endogenous controls for mRNAs and miRNAs, respectively. All primers used are listed in Table S2.

### Primary neuronal culture, in vitro electroporation and quantification of in vitro axon growth

Electroporation and culture of dissociated adult mouse DRG neurons were performed based on procedures described previously (Hur et al., 2011; Saijilafu et al., 2013) with minor changes. Briefly, DRGs were first dissected out from 8-10-week-old female CF-1 mice and digested with 1 mg/ml type 1 collagenase and 5 mg/ml dispase at 37°C for 70 min, followed by 3 times of wash with MEM containing 10% fetal bovine serum (FBS) and 1x Pen Strep (PS, ThermoFisher Scientific). The digested DRGs were then dissociated into single cells with a 1 ml pipette tip in the same medium. The dissociated cells were filtered with a 100 µm cell strainer and centrifuged. The pellet was then resuspended with 100 µl electroporation buffer (mouse neuron Nucleofector^TM^ kit, Lonza) containing either RNA oligos (0.2 nmol) mixed with GFP plasmid (10µg) or GFP plasmid only. The mixture of cells, GFP plasmid and/or RNA oligos was transferred to a 2.0-mm electroporation cuvette and electroporated with the Amaxa^TM^ Nucleofector^TM^ II (Lonza). After electroporation, the cells were immediately mixed with desired volume of pre-warmed medium mentioned above and plated onto glass coverslips pre-coated with a mixture of 100 µg/ml poly-D-lysine (Sigma-Aldrich) and 10 µg/ml laminin (ThermoFisher Scientific). After the cells fully attached to the coverslips (6 hrs), the medium was totally removed to get rid of the electroporation buffer and dead cells. The cells were then cultured in MEM containing 5% FBS, 1x GlutaMAX^TM^-I (ThermoFisher Scientific), 1x PS and antimitotic reagents (20 µM 5-fluoro-2-deoxyuridine and 20 µM uridine, Sigma-Aldrich) for 3 days. No additional growth factors were added into the culture medium. The siRNAs against mouse Lin28a (L-051530-01-0005) and Lin28b (L-063393-01-0005), and mimics of let-7a (C-310503-01-0005) and let-7b (C-310504-01-0005) were all purchased from Dharmacon.

For quantification of in vitro axon growth, the cultured neurons on the coverslips were first fixed with 4% PFA in 1x PBS and washed with 1x PBS for 3 times, and then blocked in blocking buffer (2% bovine serum albumin (BSA), 0.1% Triton X-100 in 1x PBS) for 1 hr. The neurons were then sequentially immunostained with anti-βIII tubulin antibody (Tuj1, 1:1000, BioLegend 801202) and Alexa Fluor^TM^ 568 conjugated secondary antibody (1:500, ThermoFisher Scientific) for 1 hr each, both diluted in the same blocking buffer. Three times of 10-min wash with 1x PBS was performed after each antibody incubation. The coverslips were then mounted onto glass slides with Fluoroshield^TM^ histology mounting medium (Sigma-Aldrich) and fluorescent images of the neurons were captured with a CCD camera connected to an inverted fluorescence microscope controlled by the AxioVision software (Carl Zeiss). The longest axon of each neuron was manually traced and measured with the built-in “measure/curve spline” function of the AxioVision. Only.neurons with axons longer than twice the diameter of their cell bodies were included. In each independent experiment, at least 50 neurons were measured in every group.

### in vivo electroporation of adult DRG neurons and quantification of in vivo axon regeneration

The in vivo electroporation of adult mouse DRGs was performed as previously described (Saijilafu et al., 2011). Briefly, after the mice were anesthetized, a small dorsolateral laminectomy was performed on the left side to expose the left L4/5 DRGs. Plasmids (2 µg per kind) and/or RNA oligos (0.1 nmol per kind) were injected into each DRG with a pulled glass micropipette (World Precision Instruments) connected to a Picospritzer III (20-psi pressure, 6-ms duration, Parker Hannifin). Right after the injection, in vivo electroporation was performed by applying five electric pulses (35 V, 15-ms duration, 950-ms interval) using customized tweezer-like electrodes (BTX) powered by the Electro Square Porator ECM830 (BTX). The wounds were then closed and the mice were allowed 2 days to recover. Two days later, the left sciatic nerves of the mice were exposed right below pelvis and crushed with Dumont #5 forceps. The crush sites were marked with nylon epineural sutures (size 10-0). After 2 or 3 days, the mice were transcardially perfused with 1x PBS followed by ice-cold 4% PFA in 1x PBS. The sciatic nerve segments were then dissected out and post-fixed in 4% PFA overnight at 4°C. On the next day, the nerve segments were mounted onto a glass slide and covered with a coverslip and flattened before imaging. Mice without a clearly identifiable crush site or an epineural suture were excluded from data analysis. The siRNAs and mimics used were the same as those used for in vitro electroporation. The let-7 miRNA family inhibitor (450006) was purchased from Exiqon (now YFI0450006 from Qiagen).The fluorescent siRNA control (SIC003) was purchased from Sigma-Aldrich.

For quantification of in vivo axon regeneration, fluorescent images of the whole-mount nerve segments were first acquired with the same inverted fluorescent microscope mentioned above using the MosaiX module of the AxioVision software. All identifiable GFP-labeled axons in each nerve segment were then manually traced from the crush site to the distal axonal tips to measure the length of axons. Only nerves with at least 10 identifiable axons were included in data analysis.

### Immunohistochemistry of DRG cryosections and quantification of transfection efficiency of in vivo electroporation

Two days after in vivo electroporation of fluorescent siRNA controls, mice were perfused and L4/5 DRGs were collected. Frozen DRG sections of 10 µm were obtained with a cryostat and warmed on a slide warmer at 37°C for 1 hr. After being blocked with 1x PBS containing 10% goat serum and 1% Triton X-100 at room temperature for 1 hr, the sections were then immunostained with anti-βIII tubulin antibody (Tuj1, 1:500, Biolegend 801202) overnight at 4°C, followed by Alexa Fluor^TM^ 488 conjugated secondary antibody (1:500, ThermoFisher Scientific) at room temperature for 1 hr. All antibodies were diluted with the blocking buffer. Four times of 15-min wash with 1x PBS containing 0.3% Triton X-100 was performed after each antibody incubation. The DRG sections were mounted with Fluoroshield^TM^ histology mounting medium (Sigma-Aldrich) and imaged with the inverted fluorescent microscope mentioned above. Four or five non-adjacent sections from each DRG were used for analysis. For each DRG, the transfection efficiency was calculated by dividing the of number of red fluorescent siRNA control^+^/Tuj1^+^ cells by the number of Tuj1^+^ cells.

### Intravitreal injection, optic nerve crush, and RGC axon anterograde labeling

Intravitreal viral injection, optic nerve crush and RGC axon labeling were performed as described in a previous study (Park et al., 2008). Briefly, under anesthesia, 1 µl of AAV2 virus was injected into the right vitreous humors of 6–8-week old wildtype mice with glass micropipettes connected to a Picospritzer III (Parker Hannifin) (15-psi pressure, 4-ms duration). The position and direction of the injection were well-controlled to avoid lens damage. Two weeks after viral injection, the right optic nerves of the mice were exposed intraorbitally and crushed with Dumont #5 forceps (Fine Scientific Tools) for 5 s approximately 1 mm behind the optic disc. To label RGC axons in the optic nerves, 1.5 µl of cholera toxin subunit B (CTB) conjugated with Alexa Fluor^TM^ 594 (2 µg/µl, ThermoFisher Scientific) was injected into the right vitreous humors with glass micropipettes and Picospritzer III 2 days before the mice were sacrificed by transcardial perfusion. Retinas and optic nerves were dissected out and post-fixed in 4% PFA in 1x PBS overnight at 4°C and rinsed with 1x PBS for 3 times on the next day. AAV2-GFP (SL100812) was purchased from SignaGen Laboratories. AAV2-Lin28a-Flag was packaged by SignaGen Laboratories. AAV2-shPTEN was packaged by Vigene Biosciences. All virus used had titers > 1 × 10^^13^^.

### Tissue dehydration and clearing

Dehydration and clearing of optic nerves were done based on descriptions in previous studies (Erturk et al., 2012; Luo et al., 2013). Briefly, fixed optic nerves were first dehydrated in incremental concentrations of tetrahydrofuran (TFH, 50%, 70%, 80%, 100% and 100%, %v/v in distilled water, 20 min each, Sigma-Aldrich) in amber glass bottles. Incubations were performed on an orbital shaker at room temperature. Then the nerves were incubated with benzyl alcohol/benzyl benzoate (BABB, 1:2 in volume, Sigma-Aldrich) clearing solution for 20 min. The.nerves were protected from exposure to light during the whole process to reduce photo bleaching of the fluorescence.

### Imaging and quantification of RGC axon regeneration in the optic nerve

The cleared whole-mount nerves were imaged with a 20x objective on a Zeiss LSM 510 confocal microscope equipped with a motorized stage and an LSM 510 software. For each optic nerve, Z-stack function was used to acquire stacks of 1.6-µm-thick slices, while tiling function with 10% overlap between adjacent tiles was used to scan the whole nerve. Quantification of regenerating axons in the optic nerve was done as previous described (Park et al., 2008). Specifically, every 10 continuous slices were Z-projected with maximum intensity to generate a series of Z-projection images of 16-µm-thick optical sections. At every 250-µm interval from the crush site, the number of CTB-labeled axons was counted in each optical section and the width of each optical section was measured. Both numbers were used to calculate the number of axons per micrometer of nerve width, which was then averaged over all optical sections. Σad, the total number of axons extending distance d in a nerve with a radius of r, was estimated by summing over all optical sections with a thickness t (16 µm): Σad = πr^^2^^ x (average axons/µm)/t.

### Immunohistochemistry of whole-mount retinas and quantification of RGC transduction rate and survival rate

Fixed whole-mount retinas were first radially cut into a petal shape and then stained. The procedure and reagents used were the same with those used for DRG cryosections. Fluorescent images were acquired with a 20x objective on a Zeiss LSM510 confocal microscope.

For quantification of RGC transduction rate, uninjured retinas (no optic nerve crush) were taken from 3 mice 2 weeks after intravitreal AAV2-Lin28a-Flag injection and stained with anti-βIII tubulin (Tuj1, 1:500, BioLegend 801202) and anti-Flag (1:500, Cell Signaling Technology 14793) antibodies. Six fields under 20x objective were randomly obtained from the peripheral regions of each retina. For each mouse, RGC transduction rate was represented by the ratio of total Flag^+^/Tuj1^+^ cells to total Tuj1^+^ cells. Only cells in the ganglion cell layer of each retina were counted.

For quantification of RGC survival rate, both retinas were used from AAV2-GFP or AAV2-Lin28a-Flag injected mice 2 or 4 weeks after optic nerve crush. Three mice were used in each condition. Whole-mount retinas were stained with anti-βIII tubulin antibody (Tuj1, 1:500, BioLegend 801202). Eight fields under 20x objective were randomly taken from the peripheral regions of each retina. For each mouse, RGC survival rate was calculated by dividing the number of Tuj1^+^ cells in the injured retina by that in the uninjured retina. Only cells in the ganglion cell layer of each retina were counted.

### Immunohistochemistry of retinal cryosections and image analysis

The procedure and reagents used were the same with those used for DRG cryosections. Sections of 12-µm thick were stained with anti-βIII tubulin (Tuj1, 1:500, Biolegend 801202) and anti-pS6 Ser235/236 (1:500, Cell Signaling Technology 4858S) antibodies. Fluorescent images were acquired using the inverted fluorescent microscope mentioned above. Three to five non-adjacent retinal sections from each mouse were used for analysis. For each mouse, the percentage of pS6^+^ RGCs was calculated by dividing the number of pS6^+^/Tuj1^+^ cells by the number of Tuj1^+^ cells. Only cells in the ganglion cell layer of each retina were counted.

### Quantification of backward turning rate of regenerating RGC axons

For each optic nerve, a single Z-projection image was obtained by Z-projecting all optical slices with maximum intensity. The longest and clearest 10-30 axons were selected from each nerve and their trajectories near the axonal tip were traced. Backward turning was defined when an axonal tip was facing backwards (the angle between the final direction of the axonal tip and the anterograde longitudinal axis of the optic nerve was larger than 90 degrees). Backward turning rate was calculated by number of backward turning axons/number of total axons traced. Three mice were used in each group.

## Statistics

Statistical analyses were done with GraphPad Prism 7.0 and the significance level was set as *P* < 0.05. Data are presented as mean ± SEM unless specifically mentioned. For comparisons between two groups, if the data were normalized to the control group and presented as relative level, one sample t test with hypothetical value set as 1 was used to determine the statistical significance; otherwise regular two-tailed Student’s t test was used. For comparisons among three or more groups, one-way ANOVA followed by Tukey’s multiple comparison test was used to determine the statistical significance. The “n” indicates the number of independent experiments unless specifically stated otherwise.

---

**Figure S1.**
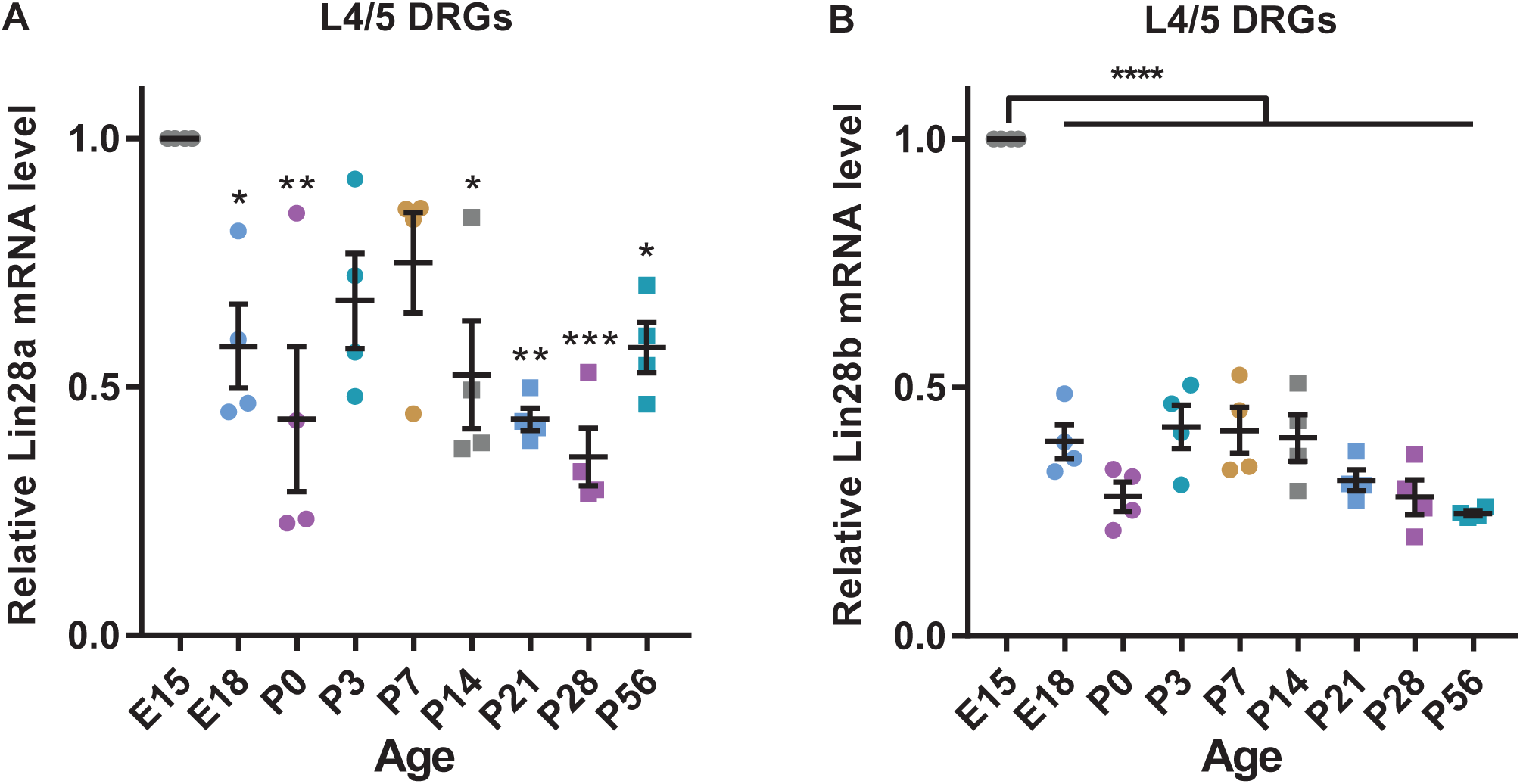
Lin28a and Lin28b mRNA levels in adult DRGs were lower than embryonic DRGs. **Related to Figure 1.** (A, B) Lin28a (A) and Lin28b (B) mRNA levels dramatically dropped from embryonic day 15 (E15) to E18 and remained at relatively lower level through postnatal day 56 (P56) (one-way ANOVA followed by Tukey’s multiple comparisons test, *P* = 0.0005 for Lin28a, *P* < 0.0001 for Lin28b, n = 4 independent experiments). \**P* < 0.05, \*\**P* < 0.01, \*\*\**P* < 0.001, \*\*\*\**P* < 0.0001, compared to control if not designated. Data are presented as mean ± SEM.

**Figure S2.**
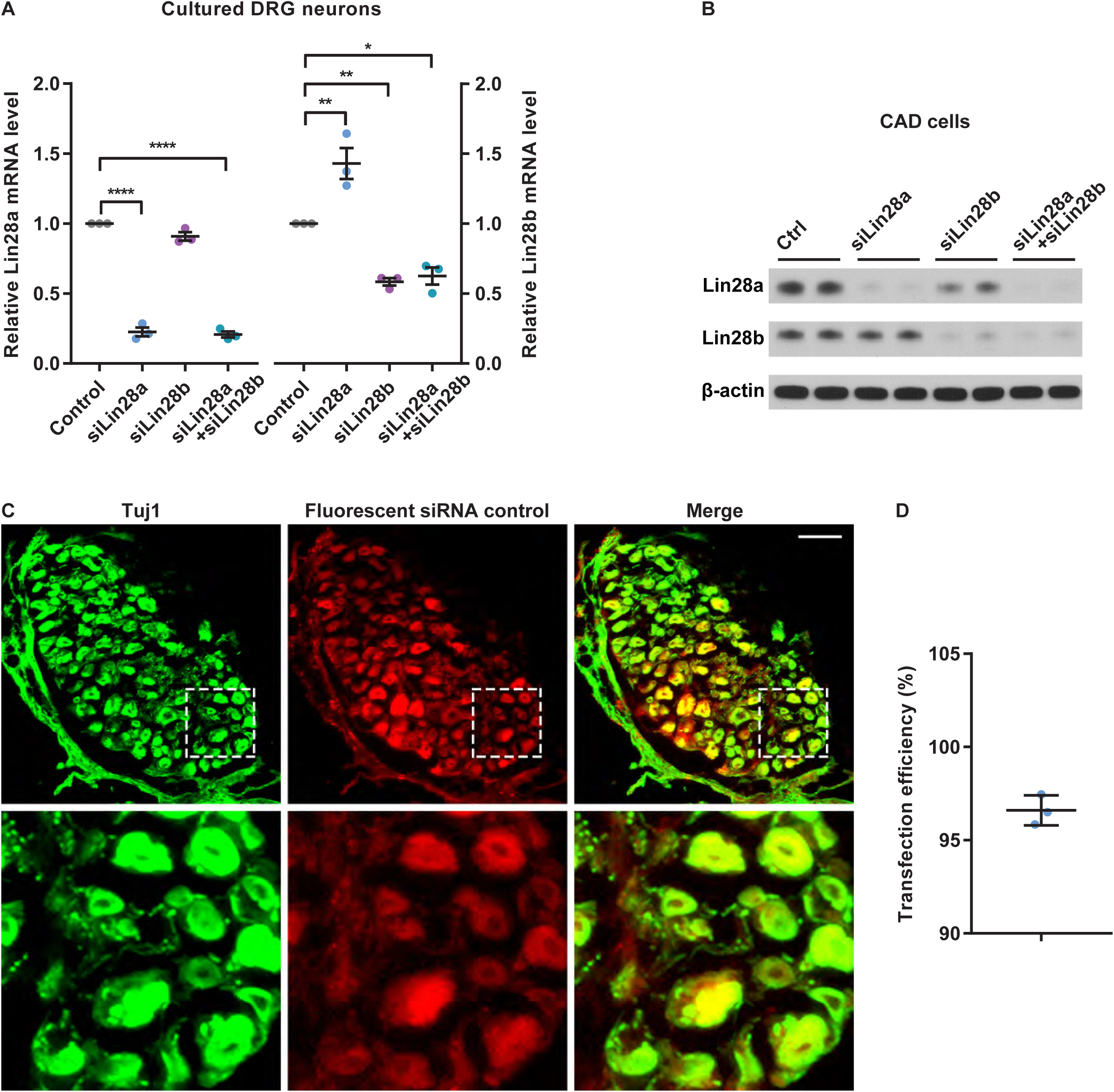
Verification of efficacy of siLin28a and siLin28b. **Related to Figure 1.** (A) Lin28a (left) and/or Lin28b (right) mRNA levels in cultured DRG neurons were lowered by Lin28a siRNA (siLin28a) and/or Lin28b siRNA (siLin28b) 3 days after electroporation (one-way ANOVA followed by Tukey’s multiple comparisons test, *P* < 0.0001 for both Lin28a and Lin28b, n = 3 independent experiments). (B) Western blot results of 2 independent experiments showing markedly diminished Lin28a and/or Lin28b protein levels in CAD cells 2 days after siLin28a and/or siLin28b transfection. (C) Representative images of DRG sections showing the high transfection efficiency of fluorescent siRNA control (red) using in vivo electroporation. DRG sections were stained with anti-βIII tubulin (Tuj1, green). The lower row shows enlarged images of areas in the dashed white boxes in the upper row. Scale bar, 100 µm for the upper row, 25 µm for the lower row. (D) Quantification of the in vivo transfection efficiency of fluorescent siRNA control in (C). The average transfection efficiency was 96.61 ± 0.8071%. Each dot represents an independently electroporated DRG (n = 3 DRGs). \**P* < 0.05, \*\**P* < 0.01, \*\*\*\**P* < 0.0001. Data are presented as mean ± SEM except in (D) where it is mean ± SD.

**Figure S3.**
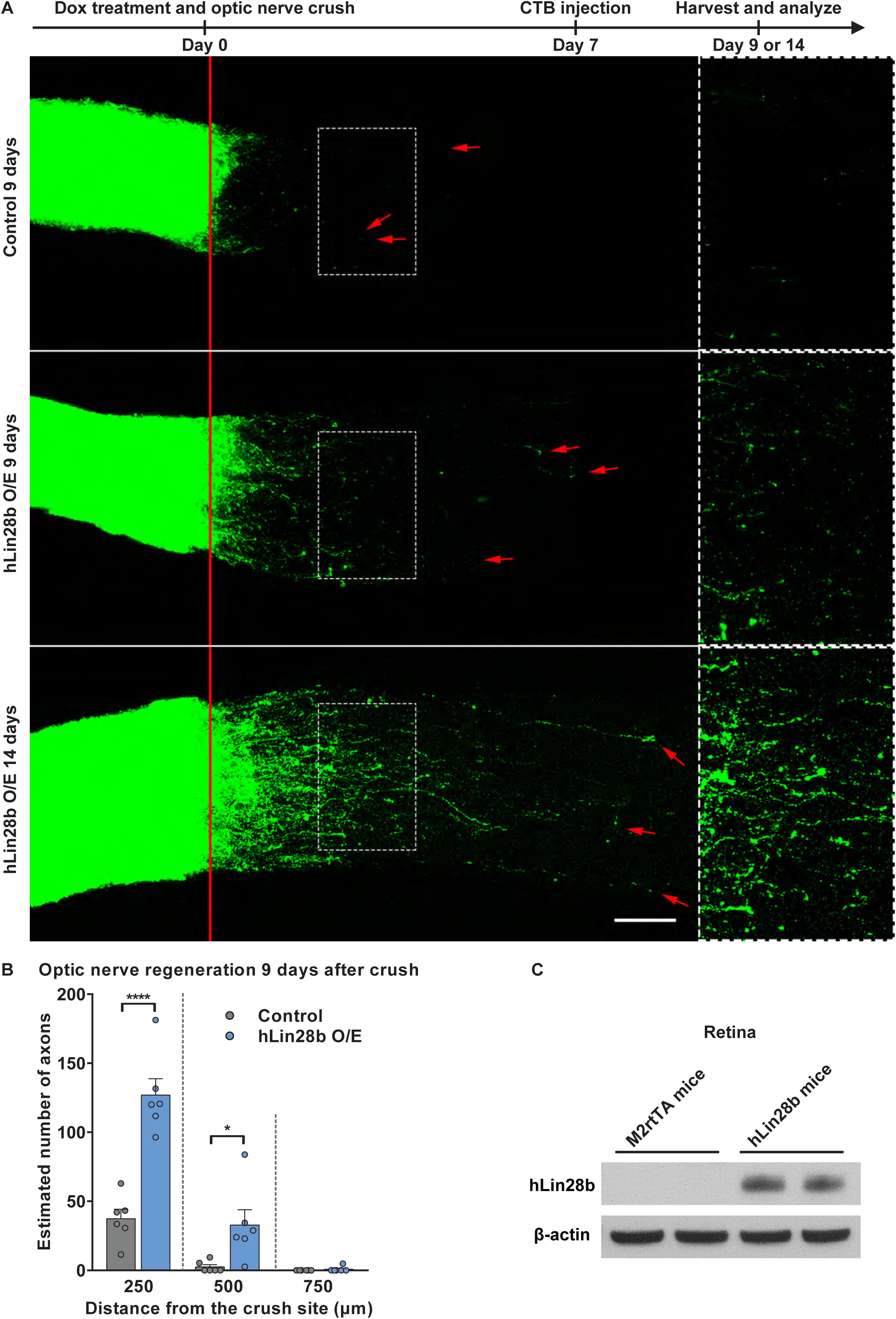
Overexpression of hLin28b in RGCs induced clear optic nerve regeneration. **Related to Figure 5.** (A) Top: diagram showing the timeline of the experiment. Bottom: representative images of cleared whole-mount optic nerves showing that overexpression of hLin28b in RGCs induced evident axon regeneration in optic nerves 9 and 14 days after nerve crush. The right column shows enlarged images of areas in the dashed white boxes in the left column. The red line indicates the crush sites. Red arrow shows the longest axon of each nerve. Scale bar, 100 µm for the left column, 50 µm for the right column. (B) Quantification of estimated number of axons at certain distances from the crush sites of optic nerves in (A) (unpaired Student’s t test, *P* < 0.0001 at 250 µm, *P* = 0.0224 and 0.3409 at 500 and 750 µm, respectively, n = 6 mice in 2-week and 4-week control groups, n = 8 mice in 2-week Lin28a overexpression group, n = 6 mice in each group). (C) Representative western blot results (2 out of 3 independent experiments) showing markedly increased hLin28b protein level in retinas of hLin28b mice 2 days after induction of hLin28b expression by doxycycline. \**P* < 0.05, \*\*\*\**P* < 0.0001. Data are presented as mean ± SEM. O/E, overexpression.

**Figure S4.**
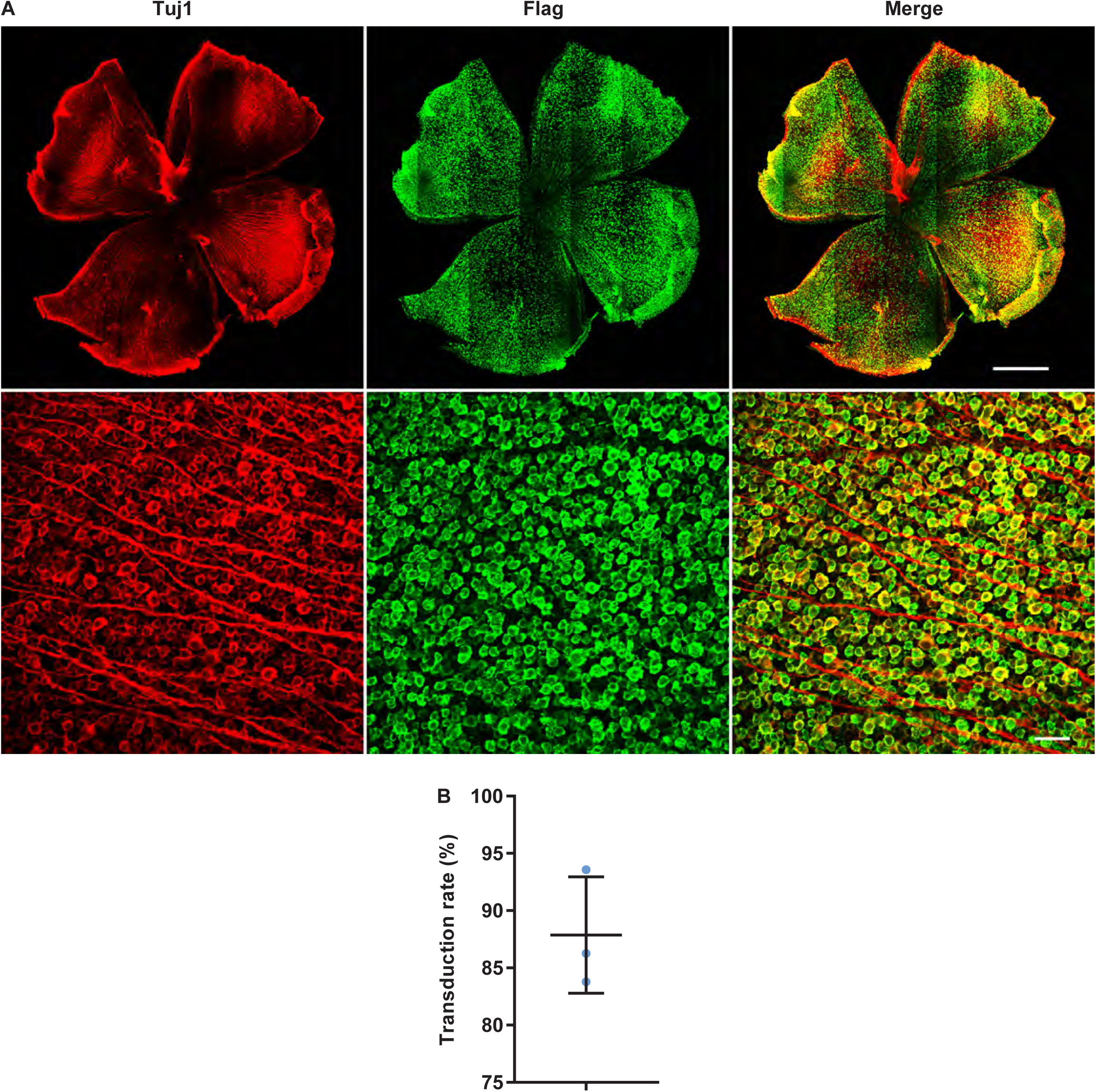
Most RGCs were transduced with AAV2-Lin28a-Flag 2 weeks after viral injection. **Related to Figure 5.** (A) Representative images of whole-mount retinas showing the high transduction rate of AAV2-Lin28a-Flag in RGCs. Whole-mount retinas were stained with anti-βIII tubulin (Tuj1, red) and anti-Flag (green) antibodies. Scale bar, 1 mm for the upper row, 50 µm for the lower row. (B) Quantification of the transduction rate of AAV2-Lin28a-Flag in RGCs in (A). The average transduction rate was 87.87 ± 5.084%. Each dot represents an independently injected retina (n = 3 retinas). Data are presented as mean ± SD.

**Figure S5.**
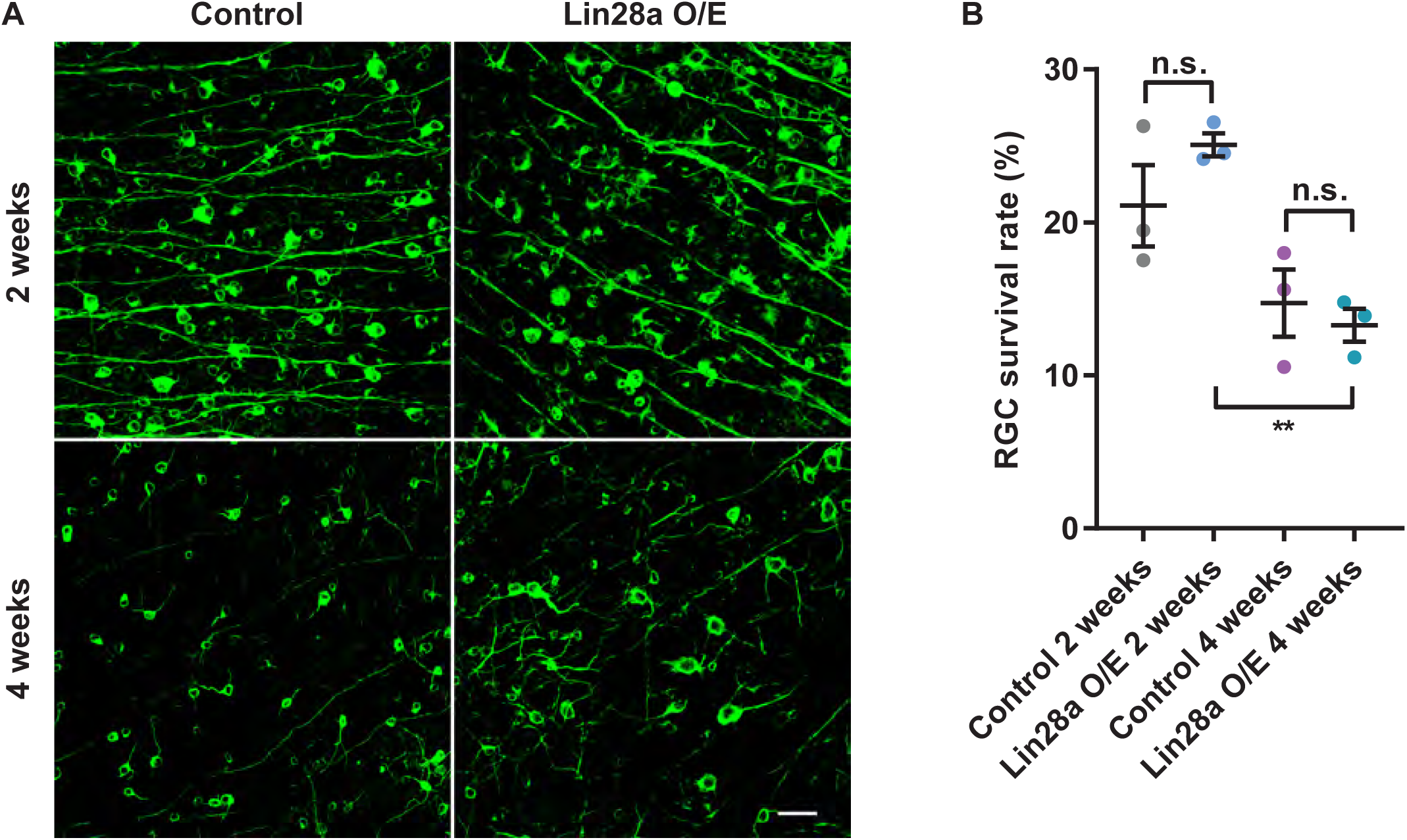
Overexpression of Lin28a did not affect survival rate of RGCs, while optic nerve crush caused continuous cell death. **Related to Figure 5.** (A) Representative images of whole-mount retinas showing that overexpression of Lin28a in retinas had no effect on RGC survival rate, while fewer RGCs survived for 4 weeks after optic nerve crush compared to 2 weeks after optic nerve crush. Whole-mount retinas were stained with anti-βIII tubulin antibody (Tuj1, green). Scale bar, 50 µm. (B) Quantification of survival rate of RGCs in (A) (one-way ANOVA followed by Tukey’s multiple comparisons test, *P* = 0.0062, n = 3 mice in each group). n.s., not significant. \*\**P* < 0.01. Data are presented as mean ± SEM. O/E, overexpression.

**Figure S6.**
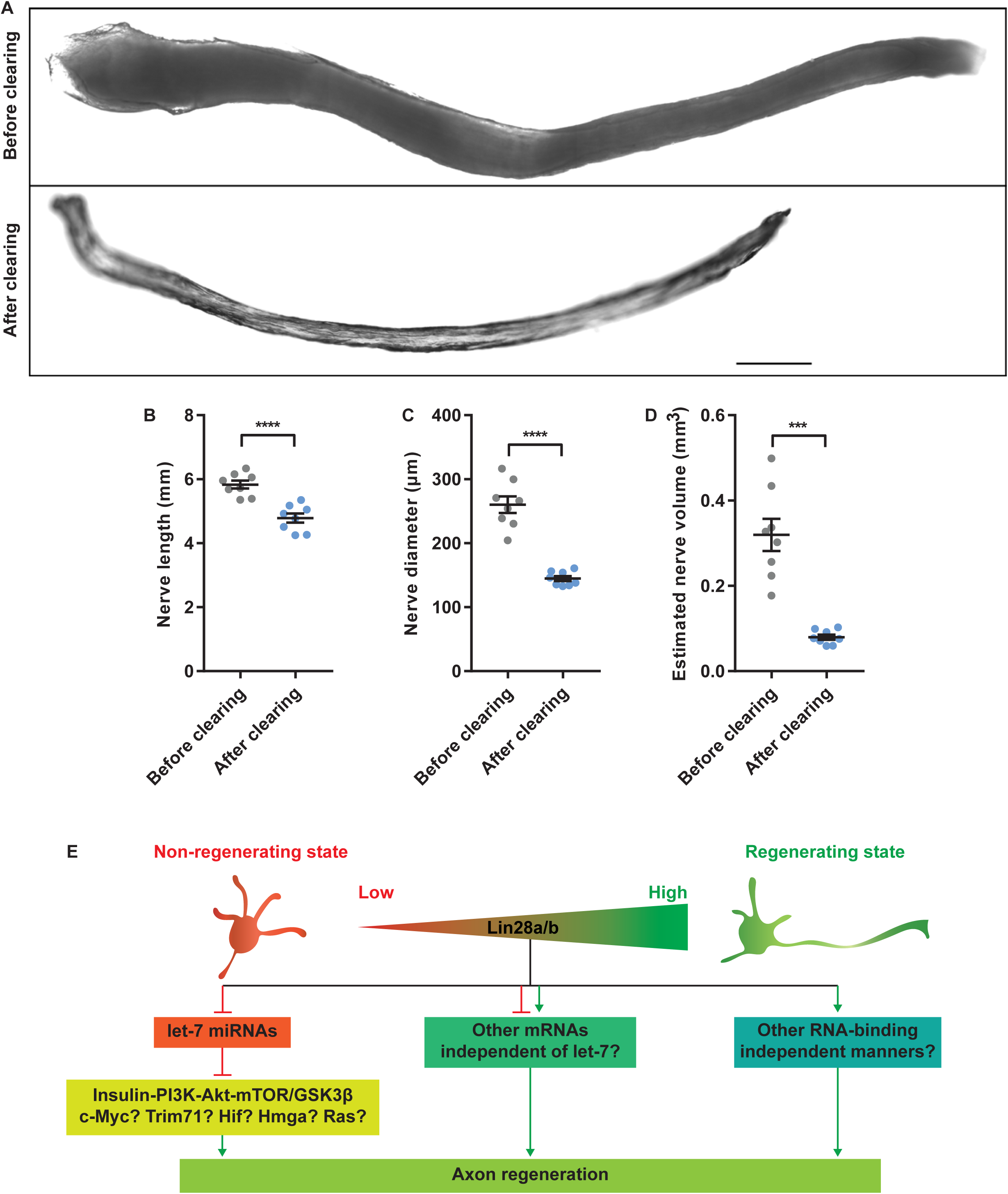
The axon length observed in optic nerves was underestimated due to tissue shrinkage caused by tissue clearing. **Related to Figure 5.** (A) Images of the same optic nerve before and after tissue clearing showing dramatic shrinkage resulting from tissue clearing. Scale bar, 500 µm. (B-D) Quantification of length (B), diameter (C) and estimated volume (D) of optic nerves before and after tissue clearing (paired Student’s t test, *P* < 0.0001 for nerve length and diameter, *P* = 0.0002 for estimated nerve volume, n = 8 optic nerves). (E) The levels of Lin28a/b are low in non-regenerating neurons, such as uninjured PNS neurons and mature CNS neurons. Upon injury, PNS neurons upregulate Lin28a/b levels to regain axon regeneration ability. Although mature CNS neurons (RGCs in our case) do not retain such intrinsic response to the injury, reintroducing Lin28a/b into mature CNS neurons might reprogram their gene expression to enhance axon regeneration. Lin28a/b upregulation represses let-7 miRNAs, leading to activation of regeneration-associated signaling pathways, such as the PI3K-Akt-mTOR/GSK3 pathways, etc. Lin28a/b may also promote axon regeneration through other mRNA targets or RNA-binding independent mechanisms. \*\*\**P* < 0.001, \*\*\*\**P* < 0.0001. Data are presented as mean ± SEM.

**Table S1.**
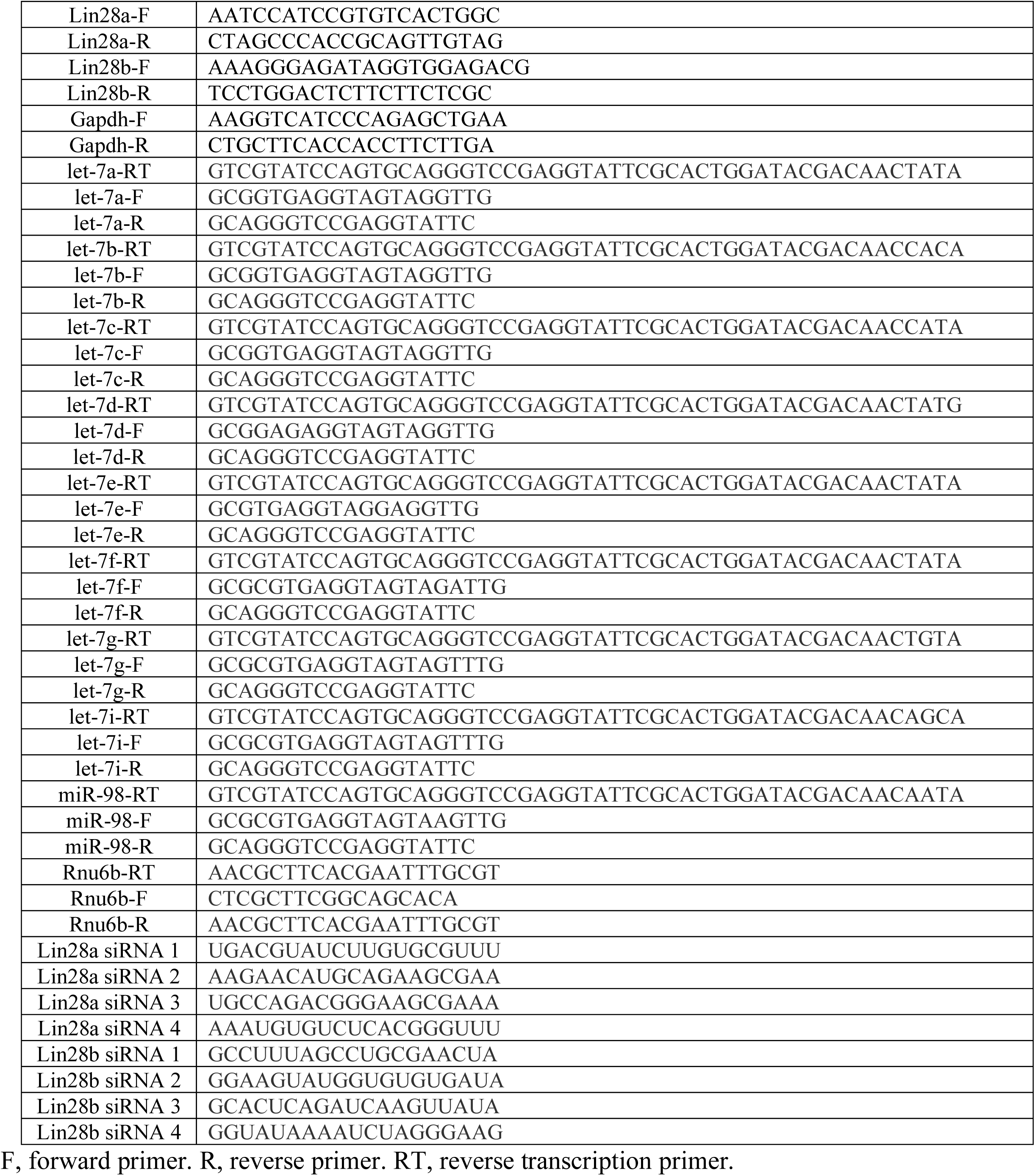
**Primers and target sequences of siRNAs used in this study. Related to Experimental Procedures.**

